# Live-cell analyses with unsegmented images to study cancer cell response to modified T cell therapy

**DOI:** 10.1101/2025.06.04.657687

**Authors:** Leo Epstein, Adam C Weiner, Archit Verma, Mozhgan Saedi, Julia Carnevale, Alex Marson, Barbara E Engelhardt

## Abstract

Live-cell imaging (LCI) of modified T cells co-cultured with cancer cells is commonly used to quantify T cell anti-cancer function. Videos captured by LCI show complex multi-cell behavioral phenotypes that go beyond simple cancer cell fluorescence measurements. Here, we develop an unsupervised analysis workflow to characterize LCI data generated using the Incucyte imaging platform. Unlike most LCI analyses, we avoid cell segmentation due to the low spatiotemporal resolution of the LCI videos and high levels of cell-cell contact. Instead, we develop methods that identify global aggregation patterns and local cellular keypoints to characterize the multicellular interactions that determine cancer cell sensitivity to, or escape from, T cell surveillance. We demonstrate our segmentation-free live-cell behavioral analysis (SF-LCBA) methods on TCR T cells from four donors with varying proportions of cells with a beneficial RASA2 knockout and effector-to-target initial concentrations in a co-culture with A375 melanoma cells. We find that different T cell modifications affect the spatiotemporal dynamics of multicellular aggregate formation. In particular, we show that fewer and smaller cancer cell aggregates form at high ratios of effector T cells to target cancer cells and high titrations of T cells with RASA2 knockouts. Our SF-LCBA method identifies, characterizes, and tracks cellular aggregate formation in datasets that are unsuitable for cell segmentation and tracking, opening the door to more therapeutically-relevant measurements of modified T cell therapy cell behavioral phenotypes from LCI data.

## Introduction

T cell cancer immunotherapies have in the past decade gained recognition for their therapeutic potential. The 2017 success of anti-CD19 chimeric antigen receptor (CAR) T cell therapy and subsequent FDA approval was a bell weather for this class of therapies for liquid cancers^1,2^. Since then, research to improve T cell therapies and adapt them for use in solid tumors has accelerated^3,4^. T cell receptor (TCR) based immunotherapies have been particularly promising; an anti-MAGE-A4 TCR T cell therapy for metastatic synovial sarcoma became the first FDA-approved T cell therapy for solid tumors in 2024^5^. TCR T cells differ from CAR T cells in that they bind to cancer-specific proteins presented on the cell surface by human leukocyte antigens (HLAs) instead of cancer cell surface proteins such as CD19. This advantage of TCR T cells increases the number of potential cancer-specific antigens that can be targeted with a T cell therapy.

Another active area of T cell cancer immunotherapy research is to improve anti-cancer activity of the modified T cell. Endogenous regulatory pathways and immunosuppressive tumor microenvironments can both trigger T cell dysfunction, which is characterized by reduced T cell proliferation and cytotoxic signaling. Fortunately, targeted gene editing with CRISPR-Cas9 has the potential to overcome these limitations and enhance therapeutic efficacy^6–10^. Several recent studies have performed genome-wide CRISPR knockout screens under *in vitro*^11–14^ and *in vivo*^14–17^ immunosuppressive conditions to identify genetic perturbations that reduce T cell therapeutic functions. These modified CAR and TCR T cells hold immense promise to improve anti-cancer cellular immunotherapy.

While genome-wide CRISPR screens successfully nominate promising T cell perturbations in a high-throughput fashion, these candidates are validated using lower throughput experiments. In particular, fluorescence-based killing assays are a common *in vitro* functional assay for quantifying anti-cancer activity^11,13,17^. In this assay, modified T cells are co-cultured with cancer cells that express a fluorescent reporter. The co-cultures are then subject to multi-channel live-cell imaging (LCI) to quantify the amount of fluorescent signal coming from cancer cells. Experimental conditions associated with the largest reduction in fluorescent intensity across the well plate, as a proxy for a reduction in viable cancer cells, are determined to have the greatest anticancer potential and then often followed-up with *in vivo* studies. However, there is often a discrepancy between *in vitro* and *in vivo* cancer cell killing efficacy^17^.

To address the discrepancy between *in vivo* and *in vitro* efficacy and to better understand the mechanisms for increased anticancer potential, computational researchers have been trying to extract additional features from co-culture live-cell imaging (LCI) data beyond cancer cell killing efficacy^18–21^. Research in this area has been made possible through critical advancements from the deep learning computer vision field to develop models that segment individual cells from phase channel images^22–27^. The resulting cell segmentation masks enable measurements of cell morphology, such as quantifying cell size and circularity, which are known to vary between cell types and phenotypic states^28–31^. In LCI datasets with high temporal resolution, cell tracking models are often run downstream of segmentation, which allows for estimation of cell motility, cell division events, and changes in cell morphology over time^32–36^. Recent work from our group has demonstrated that cell segmentation and tracking outputs can be used to compare T cell interaction times, changes in morphology, and proliferation dynamics between different genetically modified TCR T cells^31^.

Despite the promise of cell segmentation and tracking to provide informative features, these pre-trained deep learning models are often incompatible with existing cancer and T cell co-culture LCI data. Co-culture experiments test cancer cells and T cell products across many different experimental conditions and technical replicates. Given that automated LCI microscopes only have one camera per 96-well plate, there is a tradeoff between number of wells imaged and the spatiotemporal resolution of the images. The spatiotemporal resolution of co-culture experiment LCI data is commonly 5*μ*m/pixel and 2 hours between frames^11,13,17^. This is too low to perform cell segmentation and tracking for T cells, which requires ≤ 1*μ*m/pixel (≥10x magnification) and ≤ 15 minutes between frames, respectively^31,33,35^. Additionally, even when the spatiotemporal resolution criteria are satisfied, these models struggle to segment and track cells belonging to cancer cell aggregates— commonly found in co-culture LCI experiments—since cell aggregates are not present in any training data for these models. Therefore, there is a critical need to measure biologically-relevant features of co-culture LCI data, such as T cell activation and cancer cell aggregation, without the use of pre-trained cell segmentation and tracking methods.

In this paper, we analyzed LCI data of RASA2 knockout (RASA2KO) TCR T cells co-cultured with the A375 melanoma cell line^13^ to characterize cancer cell aggregation across experimental conditions. Deletion of RASA2 was chosen as the genetic pertubation for these experiments as its ablation was found to enhance MAPK signaling and T cell cytolitic activity in a genome-wide CRISPR screen. Each LCI co-culture varied in the titration of RASA2KO mutant to wildtype T cells, the effector to target (E:T) ratio, and the primary T cell donor. Since the cancer cells expressed red fluorescent protein (RFP), each well had phase and red channel images taken once every 2 hours for a total of 144 hours.

In order to circumvent low spatiotemporal imaging resolution that prevented cell segmentation and tracking of this data, we formulated a segmentation-free live-cell behavioral analysis (SF-LCBA) framework to understand the multicellular interactions associated with the observed cancer cell aggregation processes. This works builds on a segmentation-free deep learning model for classification that was naive to any observed behaviors of the cells^37^. The two main components of SF-LCBA are the calculation of cancer cell spatial entropy across each well^38^, and using scale-invariant feature transform (SIFT) to identify and characterize local keypoints in the phase channel images as one of five single or aggregate cell types^39^. SIFT extracted hundreds to thousands of keypoints per image—where each keypoint represents a single- or multi-cellular instance and is captured using a length-128 descriptor vector. We clustered SIFT descriptor vectors and interpreted the meaning of each SIFT cluster through visual inspection and comparison of RFP intensities. Statistical testing was applied to both the SIFT clusters and the spatial entropy scores to show their relationships with experimental conditions and temporal dynamics. Overall, our analysis demonstrated that cancer cell growth and, importantly, aggregation is most effectively inhibited in co-cultures with high starting doses of RASA2KO TCR T cells (high E:T ratio, high RASA2KO titration). We believe the SF-LCBA framework has broad potential to improve analyses of co-culture LCI data beyond total cancer cell counts, stratifying distinct conditions where total RFP expression would otherwise be equal in a wide variety of imaging scenarios, ultimately leading to better cellular cancer immunotherapies.

## Methods

The proof-of-concept data for our segmentation-free live-cell behavioral analyses (SF-LCBA) methods come from LCI taken with the Incucyte platform from a study of TCR T cells in four donor backgrounds. These TCR T cells with the RASA2 knockout are co-cultured with cells from the A375 malignant melanoma cell line^13^. These LCI data allow us to showcase the association of our metrics with donor, E:T ratio, and RASA2KO mutant-to-wildtype TCR T cell titration. The SF-LCBA workflow that we develop may also be applied to LCI experiments from different experimental systems as long as there are phase contrast and fluorescence channels available.

### Live-cell imaging experiments

NY-ESO1-reactive 1G4 TCR T cells were co-cultured with pre-plated red fluorescent protein positive (RFP+) A375 tumor cells in 96 well flat-bottom plates. The plates were imaged every 2 hours for 144 hours using the IncuCyte Zoom live-cell imaging platform (Essen Bioscience). Across two channels images were acquired. These channels consisted of a phase (brightfield) channel and an RFP (red) channel. The plates were laid out in a matrix as follows: Across 12 columns, T cell titration was serially diluted from 100% RASA2KO T cells to 3.1% RASA2KO T cells and 96.9% wild type T cells. These titrations were duplicated across four technical replicates. Across 14 rows, Effector T cell to cancer (A375 cancer cells) cell ratio (E:T ratio) is progressively diluted from 2.8284 by a factor of 1.14142 to 0.2. These dilutions are duplicated (across four technical replicates) for all E:T ratios except for the lowest one. Each combination of E:T ratio and RASA2KO percentage is replicated four times, except for the lowest E:T ratio, which only has two technical replicates. There are four plates, the T cells of each plate across mutational status are derived from different donors. Each experiment has approximately sixty five images; there is variation in time series length of ± 3 frames between donor replicates. The spatial resolution per image frame is 4.975 *μ*m/pixel. To ensure time frames exactly matched across conditions, we restricted our analysis of this dataset to the first 64 time frames from 5 RASA2KO titrations (100%, 50%, 25%, 12.5%, 6.25%), 5 effector to target (E:T) ratios (2.83, 2.00, 1.41, 1.00, 0.71), 4 donors, and 2 technical replicates; however, these filters are performed for simplicity of presentation and are not essential to the analyses.

### Thresholding on RFP intensity to obtain cancer cell masks

Red fluorescent protein (RFP) was primarily localized to the nucleus and cytoplasm of A375 cancer cells; however, RFP was also detected at low levels throughout the well due to background auto-fluorescence. We therefore created a binary cancer cell mask *M*(*i, j*) by thresholding the red channel’s intensity values at *≥* 3.5. We quantified the binary cancer cell mask as follows:

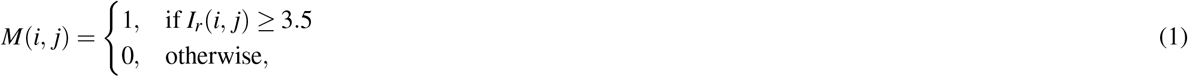

where *I*_*r*_(*i, j*) represents the red channel’s intensity value at pixel coordinates (*i, j*). *M*(*i, j*) is the resulting binary mask where *M*(*i, j*) = 1 indicates RFP+ pixels, and *M*(*i, j*) = 0 indicates RFP-pixels.

### Cancer cell mask summary statistics

The two summary statistics we computed from cancer cell masks were the cancer cell area and spatial entropy. We approximated cancer cell area for a region of interest (ROI) *R* using the ROI’s fraction of RFP+ pixels *f*_RFP+_(*R*) as follows:

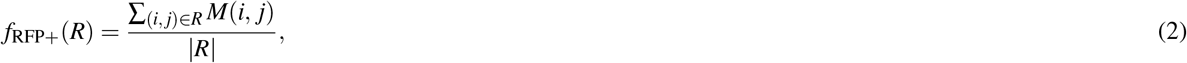

where:

- *M*(*i, j*) is the binary cancer cell mask,
- *R* is the set of pixel coordinates within the ROI, and
- |*R*| is the total number of pixels in the ROI.

To compute spatial entropy, we partitioned the binary mask *M*(*i, j*) of ROI *R* into a set of 100 × 100 pixel subregions *S*(*R*). We computed the Shannon entropy measure based on the fraction of RFP+ pixels within each region. We used the following equation to compute entropy:

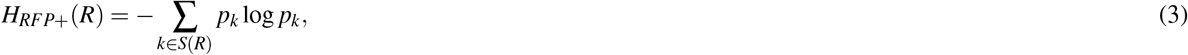

where:

- *H*_*RFP*+_(*R*) is the spatial entropy of the cancer cell mask *M*(*i, j*) in ROI *R*,
- *S*(*R*) is the set of 100 × 100 pixel subregions in ROI *R*,
- *P*_*k*_ is the proportion of RFP+ pixels in the *k*th subregion, given by 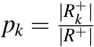 where 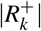 is the number of RFP+ pixels in the *k*th subregion, and |*R*^+^| is the total number of RFP+ pixels in the entire ROI *R*.

For both summary statistics, the ROI *R* can either be the entire image or a window within an image.

### Applying SIFT to LCI phase images

The scale-invariant feature transform (SIFT)^39^ was applied to all phase (brightfield) image frames in the LCI dataset using the `scikit-image` implementation with default settings of 8 octaves and 3 scales^40^. The SIFT algorithm consists of two steps: 1) identifying keypoints and 2) computing descriptors. Each SIFT keypoint is a set of image coordinates (*i, j*) that represents a local maximum or minimum of a Gaussian pyramid transform of the image. SIFT identifies keypoints across three discrete scales *σ*, and 8 octaves *o*, whereby the radius *r* of each keypoint is defined by *r* = *σ*\* 2^1+*o*^^41^. SIFT also captures the orientation angle of the gradient around every keypoint. Keypoint angle is used to standardize the orientation of the calculated descriptor histograms, insuring rotational invariance. After identifying all keypoints in an image, SIFT computes a length-128 descriptor vector for each keypoint. This vector is a flattened histogram of gradients calculated around the keypoint coordinates on the untransformed image.

### Subsampling SIFT descriptor matrix

Since SIFT key-points correspond to local maxima and minima in the feature space of a Gaussian pyramid transform of the image, we obtained *d*_*n*_ keypoints (and descriptors) for each phase image *n* instead of a constant number of keypoints per image. To limit memory usage, we randomly subsampled 10% of each image’s keypoints when constructing the original descriptor matrix of shape *D* ×128, where 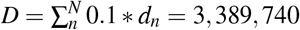. We further randomly subsampled the descriptor matrix down to *D* = 50, 000 keypoints when performing statistical tests and visualization to avoid overpowered p-values and improve interpretation, respectively.

### K-means clustering of SIFT descriptors

We used K-means to cluster the *D* × 128 SIFT descriptor matrix. We used the `scikit-learn` implementation of K-means clustering^40^ with default parameters, and we performed a parameter sweep for the number of clusters *K*, ranging from *K* = 3 to *K* = 10. We computed the silhouette score and sum of squared distances (SSD) to cluster center for each set of clustering results *K*. The silhouette score measures the quality of clustering by evaluating how similar a (key)point *i* is to its own cluster compared to other clusters. It first computes the the mean intra-cluster distance for each keypoint *a*_*i*_:

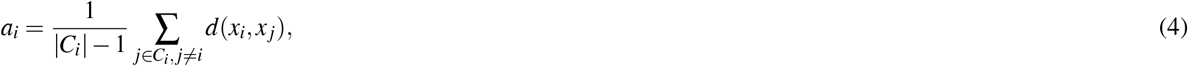

where:

- *C*_*i*_ is the cluster to which keypoint *i* belongs,
- |*C*_*i*_| is the number of keypoints in cluster *C*_*i*_,
- *x*_*i*_ and *x* _*j*_ are the length-128 descriptor vectors for the *i*th and *j*th keypoints, respectively,
- *d*(*x*_*i*_, *x* _*j*_) is the Euclidean distance between descriptor vectors *x*_*i*_ and *x* _*j*_.

The second step of computing the silhouette score is to find the nearest cluster distance *b*_*i*_, which represents the average distance of keypoint *i* to all keypoints in the nearest neighboring cluster:

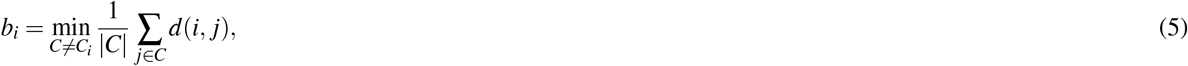

where:

- *C* is any cluster other than *C*_*i*_,
- |*C*| is the number of keypoints in cluster *C*.

Using both *b*_*i*_ and *a*_*i*_ for all keypoints, we compute the mean silhouette coefficient over all keypoints in the SIFT matrix *D*:

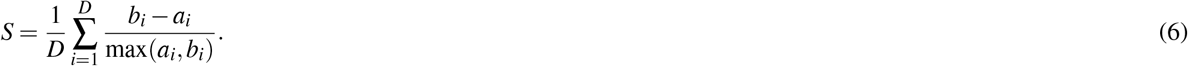

Note that we only computed silhouette score on *D* = 50, 000 keypoints since computing silhouette scores has runtime complexity of *O*(*N*^2^). Silhouette scores range from −1 to 1, where 1 represents perfectly clustered data.

The other metric we computed was the sum of squared distances (SSD) from each keypoint to the cluster center, also known as the within-cluster sum of squares (WCCS) metric:

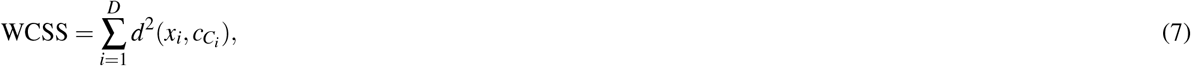

where

- *D* is the total number of keypoints,
- *x*_*i*_ is the length-128 descriptor vector for the *i*th keypoint,
- *C*_*i*_ is the cluster to which keypoint *i* belongs,
- 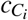 is the length-128 centroid of cluster *C*_*i*_,
- 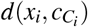 is the Euclidean distance between *x*_*i*_ and cluster center 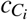.

We selected the optimal number of clusters by applying the elbow heuristic to WCCS scores and jointly examining the silhouette scores.

### Principal components analysis

We used principal component analysis (PCA) to visualize the 128-length SIFT descriptors in a lower dimensional space. To do so, we ran PCA on the full *D* × 128 descriptor matrix with *D* = 3, 389, 740 with 30 principal components (PCs). Only the first two 2 PCs (PC1 and PC2) were used in the SIFT PCA embedding plots, although we studied latter PCs for additional correlations with relevant covariates without success.

### Mixed effects linear models

We related several image-level summary statistics (cancer cell area, cancer cell spatial heterogeneity) to experimental conditions of the image (RASA2KO titration, E:T ratio, donor, technical replicate, time) using mixed effects linear models implemented in `statsmodels`^42^. Each model performed linear regression where one summary statistic was the response variable. Across models, *RASA2KO titration, E:T ratio, technical replicate*, and *time* were the fixed effect covariates, while *donor* was the random effect variable. Coefficients were estimated for each fixed effect variable, a *y*-intercept term, and all possible combinations of 2-, 3-, and 4-variable interaction terms. *Donor* and *technical replicates* were treated as categorical variables while all other variables were continuous. Categorical variables were one-hot encoded. Since we only have two technical replicates, we showed the term for replicate ID of 1 in tables and figures while replicate ID of 0 was absorbed into the intercept term. We reported the coefficients and p-values for fixed effect estimates in these models, where the p-values represent the Wald test probability of accepting the null hypothesis that a true coefficient is equal to 0. All p-values from these tests were corrected with Benjamini-Hochberg false discovery rate (FDR) correction.

### SIFT cluster enrichment testing

When testing whether certain SIFT clusters were enriched or depleted for certain experimental conditions, we had to treat categorical experimental covariates differently from continuous ones.

The categorical covariates were *donor* ID and technical replicate ID. We iterated through each cluster ID (0, 1, …, 6) and variable value (donor 0, donor 1, …, replicate 1). For each unique combination of cluster ID and variable value, we created a 2 × 2 contingency table comparing that value’s presence inside the cluster versus outside of it. The contingency tables were then used to perform *χ*-squared tests and compute the log_2_ odds ratios, yielding p-values and effect sizes, respectively.

The continuous covariates were *time, RASA2KO titration*, and *E:T ratio*. We iterated through each cluster ID and covariate type, performing a Kruskal-Wallis test to obtain a p-value and computing the Cohen’s d (standardized mean difference) effect size. Cohen’s d measures how much the mean covariate value differs between the points in a cluster and the points not in a cluster. Therefore, positive Cohen’s d means that the cluster had more points at large values for that covariate (e.g., enriched in later time frames) or fewer points at low values for that covariate (e.g., depleted in earlier time frames).

We used the subsampled set of SIFT keypoints (*D* = 50, 000) when performing these statistical tests to limit the number of p-values that were smaller than the machine float precision of 2 × 10^−308^. All p-values from these tests were corrected with Benjamini-Hochberg FDR correction.

## Results

In this paper, we examine how segmentation-free analysis of live-cell imaging (LCI) data can be used to characterize cellular behavior in cancer and T cell co-culture assays. We focused on a dataset of co-cultured RASA2KO TCR T cells and A375 cancer cells^13^. This dataset contains Incucyte images collected every two hours from wells that varied in their RASA2KO titration, effector to target (E:T) ratio, T cell donor, and technical replicate number (see Methods). We constructed a fully-observed image tensor (with no missing images) from this dataset of the following shape:

- 5 RASA2KO titrations (100%, 50%, 25%, 12.5%, 6.25%),
- 5 E:T ratios (2.83, 2.00, 1.41, 1.00, 0.71),
- 4 donors,
- 2 technical replicates,
- and 64 time frames.

Two channels—phase and red—were collected for each image as the A375 cancer cells expressed red fluorescent protein (RFP; Fig 1a). The red channel was thresholded to create a binary cancer cell mask where individual pixels were labeled RFP positive (RFP+) or RFP negative (RFP-). Qualitative assessment of cancer cell masks across E:T ratio, RASA2KO titration, and time suggested that cancer cells divided more and survived longer at low E:T ratios and low RASA2KO titrations (Fig 1b; Fig S1-S3).

**Figure 1.**
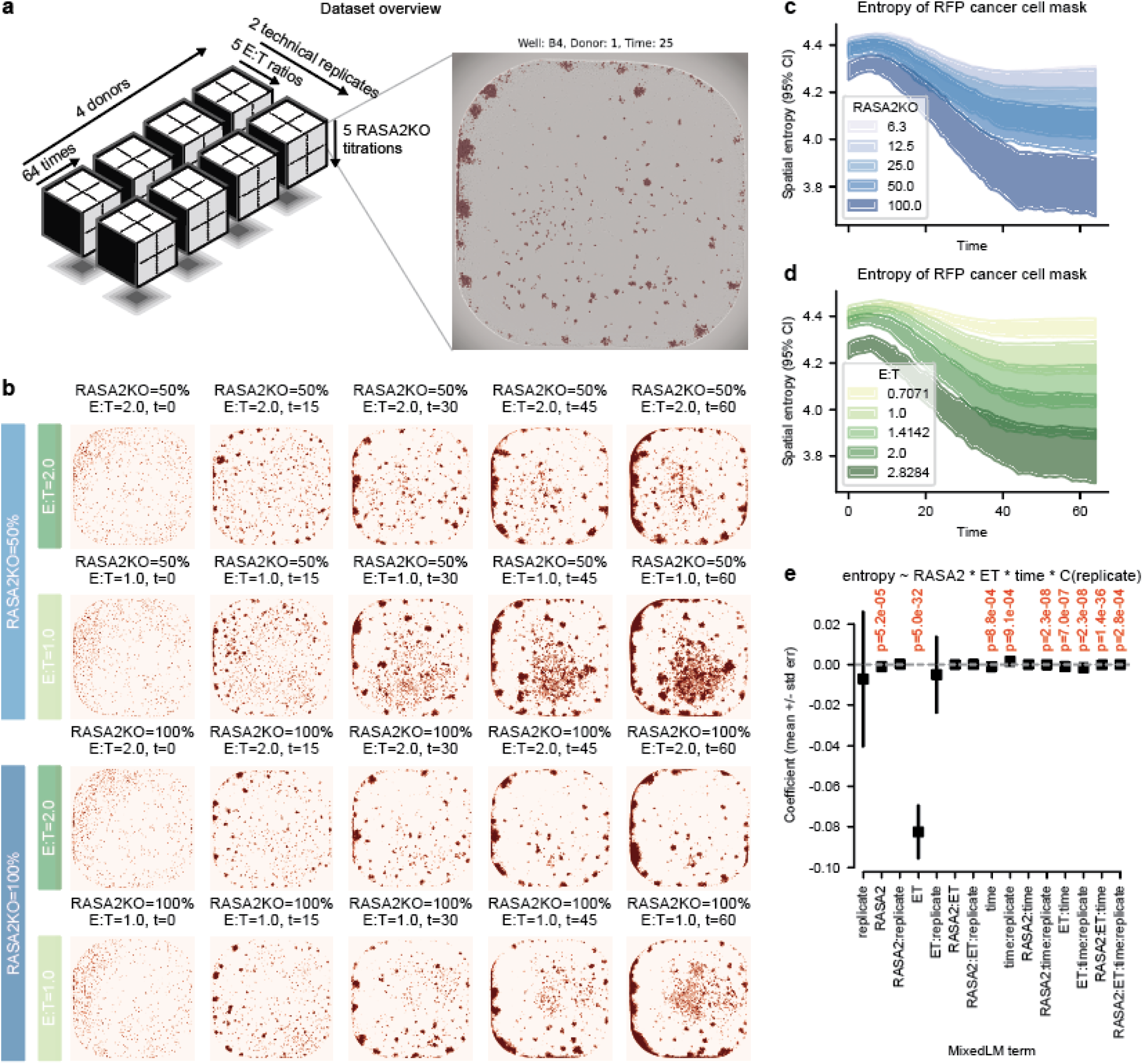
Cancer cell area and spatial entropy are computed via RFP masks. **a)** Overview of the Carnevale *et al* imaging dataset that was analyzed in this paper. 5D tensor represents the experimental metadata associated with each image. Red and brightfiend (phase) channels were captured in each image. **b)** RFP+ masks over time at varying RASA2KO titrations and E:T ratios. Time increases from left to right columns. RASA2KO titrations and E:T ratios vary between rows. **c-d)** Spatial entropy of the cancer cell (RFP+) mask over time, stratified by **c)** RASA2KO titration and **d)** E:T ratios. Shaded areas represent 95% confidence intervals. **e)** Mixed linear model results for predicting RFP spatial entropy using the covariate values for RASA2KO titration, E:T ratio, time, and technical replicate. Technical replicate ID is modeled as a categorical variable while the other covariates are modeled as continuous variables. The model is conditioned on donor ID. The intercept term is excluded here but shown in Table 2. Boxes represent mean values of each coefficient and whiskers represent +/-standard error. Interaction terms between multiple covariates are noted using colons in the x-tick labels. Adjusted P-values (Benjamini-Hochberg correction) are annotated above each term if *p*_*adj*_ < 0.05 where the null hypothesis is that the term has no effect on entropy (coefficient=0).

### Spatial entropy distinguishes cancer cell aggregation patterns across experiments

The most commonly reported metric for T cell killing assays is total cancer cell area over time^11,13,17^. Indeed, we were able to distinguish the different experimental conditions when examining the number of RFP+ pixels in each image over time (Fig S4a,b); however, solely looking at total cancer cell area did not provide insights into the spatial distribution of cancer cells, failing to suggest which cancer cells get killed by T cells or continue to proliferate. Moreover, images with identical RFP+ pixels may have wildly different spatial organizations of the cells.

**Table 1.** General linear model regression results for cancer cell mask total area. RASA2KO titration, E:T ratio, technical replicate, and time were the fixed effect variables while donor was the random effect variable. Donor and technical replicate are modeled as categorical variables and all others are continuous. Interaction terms are denoted with colons between each variable name. Adjusted p-values represent the Wald test probability of accepting the null hypothesis that a true coefficient is 0 after Benjamini-Hochberg FDR correction. All decimals are rounded to 3 digits.

|  |  |  |  |
| --- | --- | --- | --- |
| Model: | MixedLM | Dependent Variable: | RFP area |
| No. Observations: | 13446 | Method: | ML |
| No. Groups: | 4 | Scale: | 2609194480.4813 |
| Min. group size: | 3304 | Log-Likelihood: | -164861.9334 |
| Max. group size: | 3400 | Converged: | Yes |
| Mean group size: | 3361.5 |  |  |

| Term | Coef. | Std.Err. | z | $p_{adj} > \vert z\vert $ | [0.025 | 0.975] |
| --- | --- | --- | --- | --- | --- | --- |
| intercept | 22577.944 | 11473.154 | 1.968 | $9.8 \times 10^{-2}$ | 90.977 | 45064.912 |
| replicate | -9857.058 | 6141.586 | -1.605 | $1.8 \times 10^{-1}$ | -21894.345 | 2180.229 |
| RASA2KO | -8.682 | 84.119 | -0.103 | $9.2 \times 10^{-1}$ | -173.552 | 156.187 |
| RASA2KO:replicate | 210.997 | 118.962 | 1.774 | $1.4 \times 10^{-1}$ | -22.163 | 444.158 |
| ET | -513.736 | 2466.333 | -0.208 | $9.2 \times 10^{-1}$ | -5347.659 | 4320.188 |
| ET:replicate | 4867.097 | 3487.920 | 1.395 | $2.4 \times 10^{-1}$ | -1969.100 | 11703.294 |
| RASA2KO:ET | 34.040 | 47.777 | 0.712 | $6.0 \times 10^{-1}$ | -59.601 | 127.681 |
| RASA2KO:ET:replicate | -73.653 | 67.567 | -1.090 | $3.8 \times 10^{-1}$ | -206.082 | 58.775 |
| time | 7500.794 | 113.125 | 66.305 | $\leq 2 \times 10^{-308}$ | 7279.072 | 7722.516 |
| time:replicate | -17.801 | 159.983 | -0.111 | $9.2 \times 10^{-1}$ | -331.361 | 295.760 |
| RASA2KO:time | -32.328 | 2.190 | -14.760 | $1.3 \times 10^{-48}$ | -36.621 | -28.036 |
| RASA2KO:time:replicate | -18.511 | 3.098 | -5.976 | $8.6 \times 10^{-9}$ | -24.582 | -12.440 |
| ET:time | -2195.995 | 64.236 | -34.186 | $2.9 \times 10^{-255}$ | -2321.896 | -2070.095 |
| ET:time:replicate | -253.394 | 90.843 | -2.789 | $1.1 \times 10^{-2}$ | -431.444 | -75.344 |
| RASA2KO:ET:time | 6.791 | 1.244 | 5.459 | $1.4 \times 10^{-7}$ | 4.353 | 9.229 |
| RASA2KO:ET:time:replicate | 8.223 | 1.759 | 4.674 | $7.4 \times 10^{-6}$ | 4.775 | 11.671 |

**Table 2.** General linear model regression results for cancer cell mask spatial entropy. RASA2KO titration, E:T ratio, technical replicate, and time were the fixed effect variables while donor was the random effect variable. Donor and technical replicate are modeled as categorical variables and all others are continuous. Interaction terms are denoted with colons between each variable name. Adjusted p-values represent the Wald test probability of accepting the null hypothesis that a true coefficient is 0 after Benjamini-Hochberg FDR correction. All decimals are rounded to 3 digits.

|  |  |  |  |
| --- | --- | --- | --- |
| Model: | MixedLM | Dependent Variable: | RFP entropy |
| No. Observations: | 13446 | Method: | ML |
| No. Groups: | 4 | Scale: | 0.0204 |
| Min. group size: | 3304 | Log-Likelihood: | 7062.6310 |
| Max. group size: | 3400 | Converged: | Yes |
| Mean group size: | 3361.5 |  |  |

| Term | Coef. | Std.Err. | z | $p_{adj} > \vert z\vert $ | [0.025 | 0.975] |
| --- | --- | --- | --- | --- | --- | --- |
| intercept | 4.564 | 0.032 | 143.553 | $\leq 2 \times 10^{-308}$ | 4.502 | 4.626 |
| replicate | -0.007 | 0.017 | -0.415 | $7.3 \times 10^{-1}$ | -0.041 | 0.027 |
| RASA2KO | -0.001 | 0.000 | -4.256 | $5.2 \times 10^{-5}$ | -0.001 | -0.001 |
| RASA2KO:replicate | 0.000 | 0.000 | 0.497 | $7.1 \times 10^{-1}$ | -0.000 | 0.001 |
| ET | -0.082 | 0.007 | -11.948 | $5.0 \times 10^{-32}$ | -0.096 | -0.069 |
| ET:replicate | -0.005 | 0.010 | -0.508 | $7.1 \times 10^{-1}$ | -0.024 | 0.014 |
| RASA2KO:ET | 0.000 | 0.000 | 0.066 | $9.5 \times 10^{-1}$ | -0.000 | 0.000 |
| RASA2KO:ET:replicate | 0.000 | 0.000 | 0.686 | $6.7 \times 10^{-1}$ | -0.000 | 0.001 |
| time | -0.001 | 0.000 | -3.497 | $8.8 \times 10^{-4}$ | -0.002 | -0.000 |
| time:replicate | 0.002 | 0.000 | 3.457 | $9.1 \times 10^{-4}$ | 0.001 | 0.002 |
| RASA2KO:time | 0.000 | 0.000 | 1.978 | $7.2 \times 10^{-2}$ | 0.000 | 0.000 |
| RASA2KO:time:replicate | -0.000 | 0.000 | -5.838 | $2.3 \times 10^{-8}$ | -0.000 | -0.000 |
| ET:time | -0.001 | 0.000 | -5.170 | $7.0 \times 10^{-7}$ | -0.001 | -0.001 |
| ET:time:replicate | -0.001 | 0.000 | -5.810 | $2.3 \times 10^{-8}$ | -0.002 | -0.001 |
| RASA2KO:ET:time | -0.000 | 0.000 | -12.846 | $1.4 \times 10^{-36}$ | -0.000 | -0.000 |
| RASA2KO:ET:time:replicate | 0.000 | 0.000 | 3.829 | $2.8 \times 10^{-4}$ | 0.000 | 0.000 |

We were interested in developing a metric to quantify different aggregation dynamics in live-cell imaging experiments. To do this, we first decided to calculate spatial entropy^38^, which measures how evenly cancer cells are dispersed across the well. High entropy represents uniform cellular distributions and low entropy represents uneven distributions in 2D space. We found that the spatial entropy across RASA2KO titration (Fig 1c) and E:T ratio (Fig 1d) showed distinctive time-indexed curves in which entropy decreased over time, indicating that cancer cells become more less uniformly distributed across the well as they evade T cell surveillance or become crippled through T cell interactions. These curves also revealed that wells with high E:T ratio or high RASA2KO titration underwent the largest decrease in spatial entropy over time. This result suggests that, while high E:T ratio and high RASA2KO titration led to the largest reduction in cancer cell area, certain areas of each well had more dramatic reduction of cancer cells than others.

### Linear mixed model relates experimental conditions to spatial metrics

We next wanted to quantify the relative contribution of each experimental condition to the spatial metrics of cancer cell area and entropy. We fit two mixed-effects linear models, one that predicts *area* and one that predicts *entropy*, using covariates of *RASA2KO titration, E:T ratio, time*, and *technical replicate ID* (see Methods). When modeling cancer cell area, we observed area increased over time across all conditions (*p*_*adj*_ ≤ 2 × 10^−308^; Fig S4c; Table 1). While the terms for E:T ratio and RASA2KO were not significant on their own, the interaction terms with time revealed that cancer cell area increased faster at low RASA2KO titrations (*p*_*adj*_ ≤ 1.3 × 10^−48^) and low E:T ratios (*p*_*adj*_ ≤ 2.9 × 10^−255^).

When modeling spatial entropy, the linear mixed-effects model revealed that cancer cell entropy was lower at high RASA2KO titrations (*p*_*adj*_ ≤ 5.2 × 10^−5^), high E:T ratios (*p*_*adj*_ ≤ 5.0 × 10^−32^), and later time frames (*p*_*adj*_ ≤8.8 × 10^−4^; Fig 1c-e; Table 2). The fact that the RASA2KO and E:T solo terms showed greater significance than their interaction terms with time (*p*_*adj*_ ≤ 7.2 × 10^−2^ and *p*_*adj*_ ≤ 7.0 × 10^−7^, respectively) suggests that these experimental T cell parameters influence spatial entropy across all frames – not only in a time-dependent manner. Another significant term in the spatial entropy model was the three-way interaction term between RASA2KO, E:T, and time (*p*_*adj*_ ≤ 1.4 × 10^−36^), finding that cellular entropy was lowest when all three terms were high together. Together, these cancer cell mask statistics show that higher doses of RASA2KO T cells lead to less cancer cell growth and less uniform spatial configurations of cancer cells.

### SIFT feature detection and descriptor vectors from phase images

While computing spatial statistics on the RFP cell masks showed that high concentrations of RASA2KO T cells led to reduction in both cancer cell area and spatial entropy, these statistics failed to quantify the number of cancer cells that appear as singlets or in multi-cellular aggregates between different RASA2KO T cell concentrations. If cancer aggregates were larger and more frequent at high RASA2KO T cell concentrations, this would indicate that aggregates mostly consist of debris clumps or damaged cancer cells that migrate towards one another after sublethal cytotoxicity^43^. Conversely, if cancer aggregates were smaller and less frequent at high RASA2KO T cell concentrations, it would indicate that they mostly consist of proliferating cancer cells that evade T cell surveillance.

Additionally, computing spatial statistics on cell masks can only be applied to cells that express a fluorescent reporter. In the context of the RASA2KO LCI dataset^13^, we cannot examine how T cells grow, move, and change shape over time since they are not fluorescently labeled. Being limited to analysis of fluorescent channels would be particularly problematic for next-generation co-culture experiments where the number of immune cell subtypes (e.g., monocytes, B cells, dendritic cells) exceeds the number of available fluorescent channels^44^ or patient-derived cancer cells lack a fluorescent reporter^45^. For analysis of the raw phase images, one could pass all images through an autoencoder^46,47^to create a low-dimensional embedding; however, each point in the resulting embedding would represent an entire image, and not each of the hundreds of cells within that image, preventing more granular, localized downstream analysis of cellular behavior (Fig S5). We therefore wished to explore analyses that could extract single- and multi-cell instances such as cancer cell singlets, cancer cell aggregates, and cancer-T cell interactions within individual phase images without using segmentation methods. Such analyses would allow us to tease apart the nuanced cellular interactions and aggregation dynamics that are unobservable from image-level statistics.

We chose to use the scale-invariant feature transform (SIFT) algorithm^39^ to analyze the phase images. For each image in our dataset *N*, SIFT extracts *d*_*n*_ keypoints per image and computes a length-128 descriptor vector that encodes image properties surrounding each keypoint (Fig 2a, left). We then subsample the number of keypoints and concatenate across all images, producing a SIFT descriptor matrix with *D* (total number of descriptors) rows and 128 columns (Fig 2a, right; see Methods for full details). In our dataset, there were 2590 +/-1528 (mean +/-stdev) keypoints per image (Fig 2b). When cross-referencing the number of keypoints per image to the total cancer cell area obtained from the corresponding RFP mask, we found a positive correlation (Pearson *r* = 0.93, *p* ≤ 2 × 10^−308^), suggesting that the number of keypoints per image roughly approximates the cell count (Fig 2c). The original SIFT descriptor matrix had *D* ≈ 35 million descriptors across *N* = 13, 446 images; however, for computational expediency, we only included 10% of each image’s descriptors in the final SIFT descriptor matrix. The analyses proceeds identically when all of the keypoints are included.

**Figure 2.**
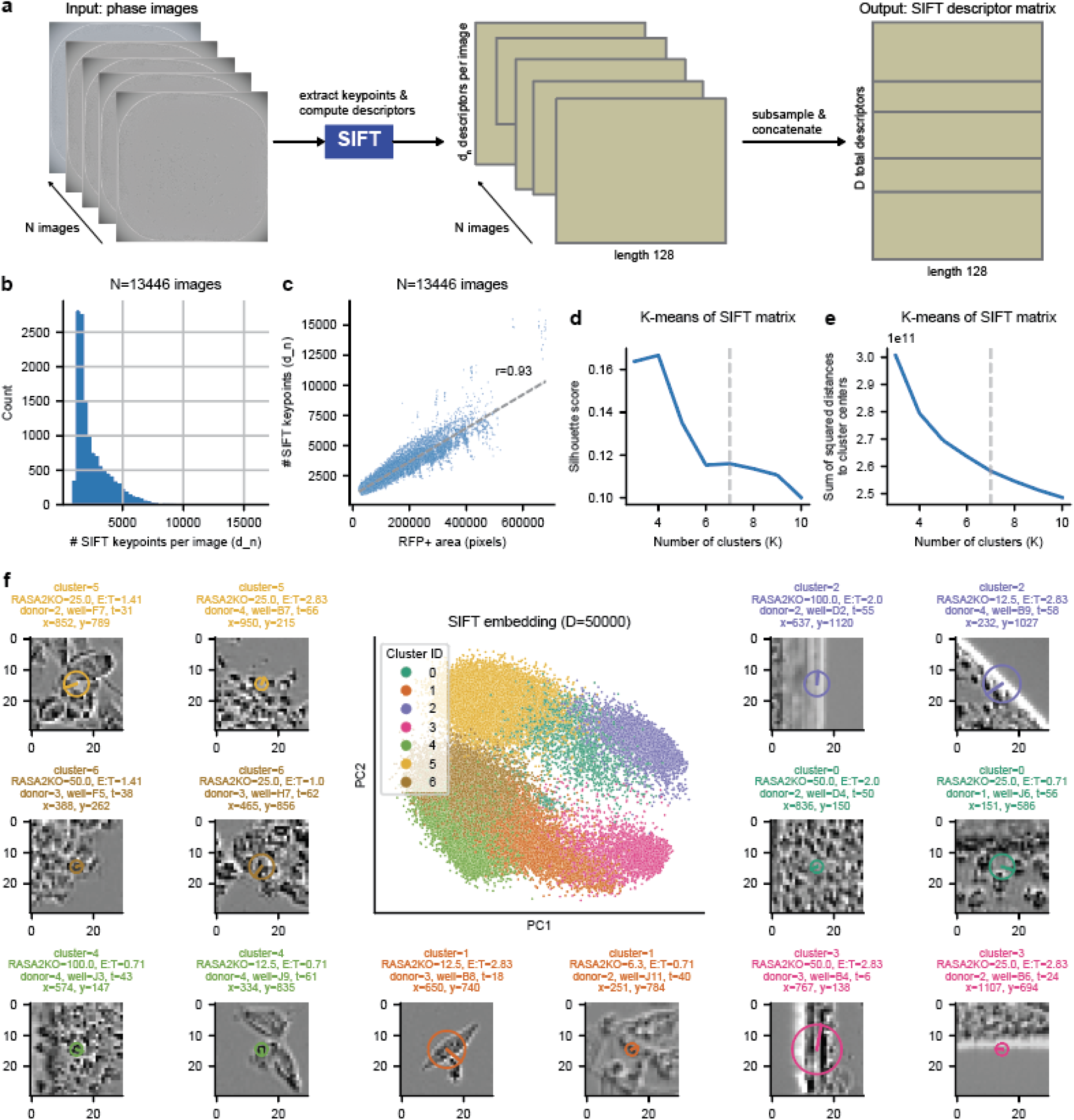
SIFT extracts relevant keypoints and computes their corresponding descriptor vectors from phase images. **a)** Schematic diagram for extracting matrix of SIFT descriptors from all phase images. Each image is passed through SIFT independently to obtain *d*_*n*_ SIFT descriptors for each image *n*. We then concatenate the SIFT descriptor matrices across all *N* images to obtain a final SIFT descriptor matrix of shape *D* × 128. We commonly only use a subset of all rows in this SIFT descriptor matrix to speed up computational analysis. **b)** Histogram of the total number of SIFT keypoints extracted per image (*d*_*n*_) prior to any subsampling. Distribution is shown for all *N* = 13446 images analyzed. **c)** Relationship between total number of SIFT keypoints and the cancer cell RFP+ area of each image. Relationship is shown for all *N* = 13446 images analyzed. The linear regression line of best fit is annotated with a dashed gray line alongside the Pearson correlation *r* value. **d)** K-means clustering silhouette scores for different numbers of clusters K. K-means clustering was performed on all *D* = 3, 389, 740 SIFT descriptors but the Silhouette scores were computed on a *D* = 50, 000 subset. **e)** Sum of squared distances from each point to its K-means cluster center for different number of clusters K. K-means clustering and the sum of squared distance calculation were both performed on all *D* = 3, 389, 740 SIFT descriptors. **f)** PCA embedding of *D* = 50, 000 SIFT descriptors colored by their K-means cluster ID. PCA was run on all *D* = 3, 389, 740 SIFT descriptors prior to subsetting to *D* = 50, 000. Two representative ROI phase images are shown for each cluster. The circle in each ROI represents the radius of each SIFT keypoint and the line represents its relative orientation. ROIs are annotated with the SIFT keypoint’s xy-coordinate along with the associated image metadata.

We next wanted to cluster the SIFT descriptor matrix to group keypoints by their similarity to one another. We performed K-means clustering of the SIFT descriptor matrix, performing a parameter sweep over the number of clusters *K* to find the optimal cluster size. For each clustering *K*, we examined the silhouette score, the sum of squared distances to cluster centers, and the cluster IDs annotated onto an embedding of the first two principal components (Fig 2d,e; Fig S6a-h). Evaluating these three components led us to select *K* = 7 for downstream analysis (see Methods). The first two principal components cumulatively explained only 21% of the total variance in the SIFT descriptor matrix (Fig S6i-n), indicating that the clusters capture additional complexity beyond what is shown in the PC1 versus PC2 SIFT embeddings.

### SIFT cluster characterization

We next define and give a label to the image properties that united keypoints within a SIFT cluster. We started by visually inspecting SIFT keypoint 30 × 30 pixel regions of interest (ROIs), both with and without RFP masks, and investigated 400 × 400 pixel ROIs containing dozens of keypoints per ROI (Fig 2f; Fig S7; Fig S8). We made the following observations about each cluster from visual inspection:

- Clusters 2 and 3 were “edge” clusters as they were located near the polystyrene edges of each well;
- Clusters 5 and 6 were “aggregate” clusters with high cell density;
- Clusters 1 and 4 were “singlet” clusters as the ROIs were commonly centered around a single, often polar, cell rather than a group of cells;
- Cluster 0 was a mixture of “aggregates” and “singlets” with a subset of this cluster representing well-wide polystyrene reflections.

We next performed a series of analyses to quantitatively support these cluster definitions.

We first wished to quantitatively support our observation that keypoints in clusters 2, 3, and a subset of 0 captured properties of the polystyrene well rather than the cells inside. To support the observation that clusters 2 and 3 were truly at well edges, we computed the distance of each keypoint to the center of the well. We found that distance to the center of the well was substantially larger in edge effect clusters compared to all other clusters (independent t-test *p* ≤ 2 × 10^−308^, *t* = − 227.3) with the standard deviation in distance being 3.8-fold smaller in edge clusters (stdev of 37 in edge clusters, 142 in other clusters; Fig 3a; Fig S9a). To determine which keypoints represented well-wide polystyrene reflections, we examined the radius (*r*) of each keypoint. We found that 3.4% of all keypoints belonging to cluster 0 had an ROI radius > 250 pixels (72 / 2109) while the largest ROI radius outside of cluster 0 was 24 pixels, indicating that the well-wide polystyrene reflections were exclusive to a subset of cluster 0.

**Figure 3.**
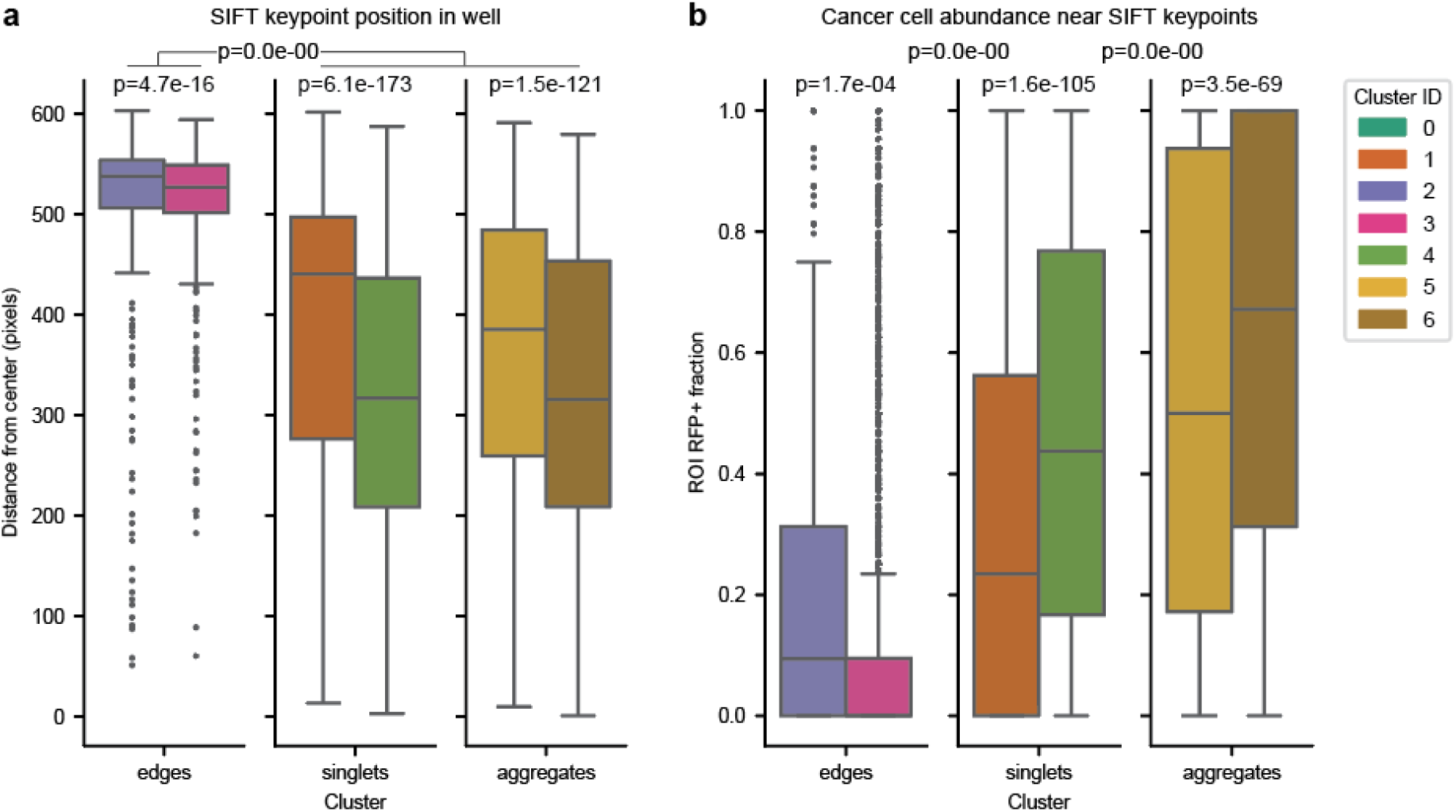
Image properties of the SIFT clusters. **a)** Distance of SIFT keypoints from the center of the well and **b)** fraction of RFP+ pixels in keypoint ROIs, grouped by cluster ID and cluster group. Edges consist of cluster IDs 2 & 3, singlets of cluster IDs 1 & 4, and aggregates of cluster IDs 5 & 6. All p-values represent independent t-tests. P-values within each cluster group compare its two cluster IDs against one another. Other p-values compare the edges group vs non-edges group in panel **a** and compare adjacent cluster groups to one another in panel **b**. Boxes represent the quartiles of the dataset while the whiskers extend up to 1.5x the interquartile range. Outlier points beyond the whiskers are drawn with small circles.

We next wished to more clearly define the singlet and aggregate clusters from one another. We hypothesized that aggregate clusters should have higher cancer cell density than singlet clusters. We therefore computed the fraction of RFP+ pixels within each keypoint’s ROI and compared these values between clusters. We found that aggregate clusters 5 and 6 had a substantially higher fraction of RFP+ pixels than singlet clusters 1 and 4 (independent t-test *p* ≤ 2 × 10^−308^; *t* = 38.5), confirming our qualitative observation that there is higher cancer cell density in aggregate keypoints compared to singlet keypoints (Fig 3b; Fig S9b).

We also compared the RFP+ fraction of ROIs within cluster types. Between the two singlet clusters, cluster 4 had a higher fraction of RFP+ pixels than cluster 1 (independent t-test *p* ≤ 1.6 × 10^−105^; *t* = 22.1; Fig 3b). Upon further exploration of keypoint ROIs, we believe this relationship reflects that cluster 1 has more T cell singlets than cluster 4 and that cluster 4 singlets contain neighboring cancer cells more often than cluster 1. Between the two aggregate clusters, cluster 6 had a substantially higher fraction of RFP+ pixels than cluster 5 (independent t-test *p* ≤ 3.5 × 10^−69^; *t* = 17.7; Fig 3b). This relationship suggests that cluster 6 aggregates have a higher cancer cell density than cluster 5. A summary of all cluster labels is shown in Table 3.

**Table 3.** Summary of SIFT clusters. Cellular clusters can consist of either singlets (single cells) or aggregates (multiple cells). Edge effect clusters were located at the edges of each well and therefore were not centered on particular cells. Cancer cell abundace was determined based on fraction of RFP+ pixels in ROIs.

| SIFT cluster ID | Name | Group | Group type | Cancer cell abundance |
| --- | --- | --- | --- | --- |
| 0 | well reflections | mixed | mixed | low-medium |
| 1 | cancer-poor singlets | singlet | cellular | medium |
| 2 | edge 1 | edge effect | well | low |
| 3 | edge 2 | edge effect | well | low |
| 4 | cancer-rich singlets | singlet | cellular | medium-high |
| 5 | cancer-poor aggregates | aggregate | cellular | medium-high |
| 6 | cancer-rich aggregates | aggregate | cellular | high |

### Enrichment of SIFT clusters between experimental conditions

We identified clusters that were enriched for certain experimental conditions over others. Such statistical testing is critical for quantifying how experimental perturbations alter the abundance of different cellular states captured by our phase images. In this dataset, we tested for cluster enrichment against the experiment variables of technical replicate ID, donor ID, time, RASA2KO titration, and E:T ratio (Fig 4a-f). We treated time, RASA2KO titration, and E:T ratio as continuous variables while donor ID and technical replicate ID were treated as categorical for statistical testing (see Methods). After false-discovery correction, no cluster was enriched or depleted beyond *p*_*adj*_ < 10^−50^ for a specific donor or technical replicate, indicating that phase images were largely similar across donors and technical replicates. However, there were nine cluster × covariate interactions across time, E:T ratio, and RASA2KO titration covariates showing substantial cluster enrichment (*p*_*adj*_ < 10^−50^; Fig 4g,h). A table of all p-values and effect sizes can be found in Supplementary File 1.

**Figure 4.**
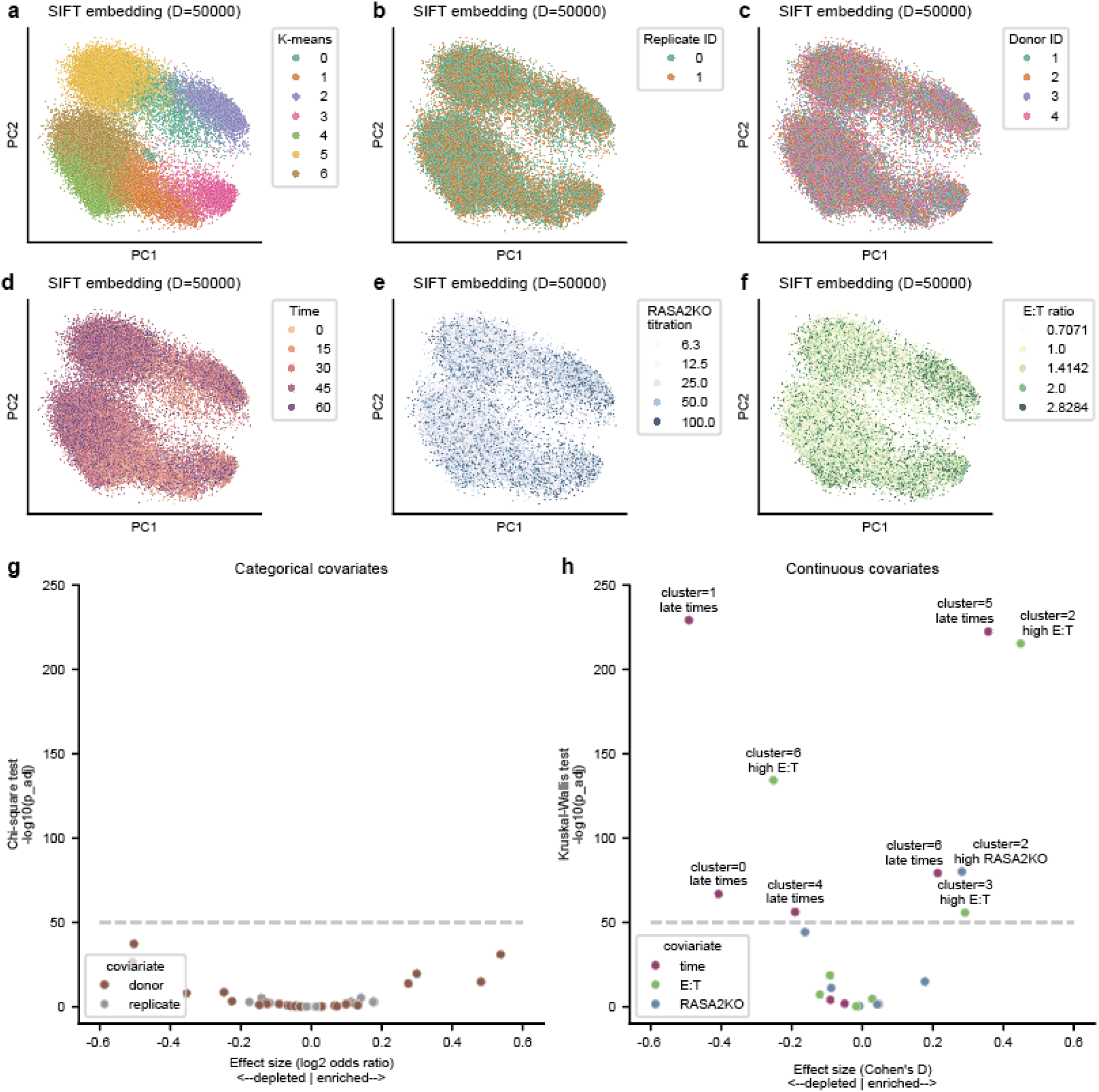
Enrichment of SIFT clusters between experimental conditions. **a-f)** PCA embedding of *D* = 50, 000 SIFT descriptors colored by **a)** K-means cluster ID, **b)** technical replicate ID, **c)** donor ID, **d)** time frame, **e)** RASA2KO titrations, and **f)** E:T ratios. **g-h)** Volcano plots testing for enrichment and depletion of covariate values within clusters. The y-axis is − log_10_ adjusted p-values and x-axis is effect size. Cluster × covariate value combinations are annotated with the color of the covariate and those with *p*_*adj*_ < 10^−50^ are also annotated with the cluster ID and covariate value. Benjamini-Hochberg correction is applied to all p-values. **g)** A chi-squared test and log_2_ odds ratio effect size are used for the covariates treated as categorical variables (donor ID and technical replicate ID). **h)** A Kruskal-Wallis test and Cohen’s d effect size are used for continuous variables (E:T ratio, RASA2KO, and time). Positive Cohen’s d for time means the cluster is enriched at late time points, high E:T ratios, or high RASA2KO titrations.

Looking into these enrichments more carefully, we found that edge effect clusters (2 and 3) were enriched at high E:T ratios and high RASA2KO titrations (*p*_*adj*_ ≤ 5.0 × 10^−216^, Cohen’s d = 0.45 for cluster 2 × E:T; *p*_*adj*_ ≤ 6.9 × 10^−81^, Cohen’s d = 0.28 for cluster 2 × RASA2KO; *p*_*adj*_ ≤ 1.6 × 10^−56^, Cohen’s d = 0.29 for cluster 3 × E:T; *p*_*adj*_ ≤ 1.1 × 10^−15^, Cohen’s d = 0.18 for cluster 3 × E:T), likely because there are few cells in the center of the well when T cells successfully limit cancer cell growth. Interestingly, both singlet clusters (1 and 4) were enriched at early time frames (*p*_*adj*_ ≤ 6.1 × 10^−230^, Cohen’s d = − 0.49 for cluster 1× time; *p*_*adj*_ ≤ 5.8 × 10^−57^, Cohen’s d = − 0.19 for cluster 4 × time) but were not enriched for any RASA2KO titrations and E:T ratios. These results suggest that singlets disappear over time regardless of whether their absence is caused by 1) increased T cell killing of cancer cell singlets at high RASA2KO T cell concentrations, or 2) unconstrained cancer cell proliferation that turns singlets into aggregates at low RASA2KO T cell concentrations. Finally, we found that both aggregate cell clusters (5 and 6) were enriched at late time frames (*p* ≤ 3.7 × 10^−223^, Cohen’s d = 0.36 for cluster 5 × time; *p* ≤ 5.2 × 10^−80^, Cohen’s d = 0.21 for cluster 6 × time), depleted at high E:T ratios (*p* ≤ 2.3 ×10^−19^, Cohen’s d =− 0.09 for cluster 5 × E:T; *p* ≤ 5.6 × 10^−135^, Cohen’s d = −0.25 for cluster 6 × E:T), and depleted at high RASA2KO titrations (*p* ≤ 7.5 × 10^−12^, Cohen’s d = − 0.09 for cluster 5 × RASA2KO; *p* ≤ 5.7 × 10^−45^, Cohen’s d = − 0.16 for cluster 6 × RASA2KO). The depletion of aggregates in wells with high RASA2KO T cell concentrations shows that the presence of these modified T cells limits the absolute number of cancer cells in aggregates. Moreover, these findings suggest that the lower entropy at high RASA2KO T cell concentrations (Fig 1) were a consequence of anticancer T cell activity creating more empty space between cancer aggregates in the RFP masks rather than more cancer cells belonging to aggregates. A graphical summary of these findings and their relationship to underlying cellular dynamics is illustrated in Fig 5.

**Figure 5.**
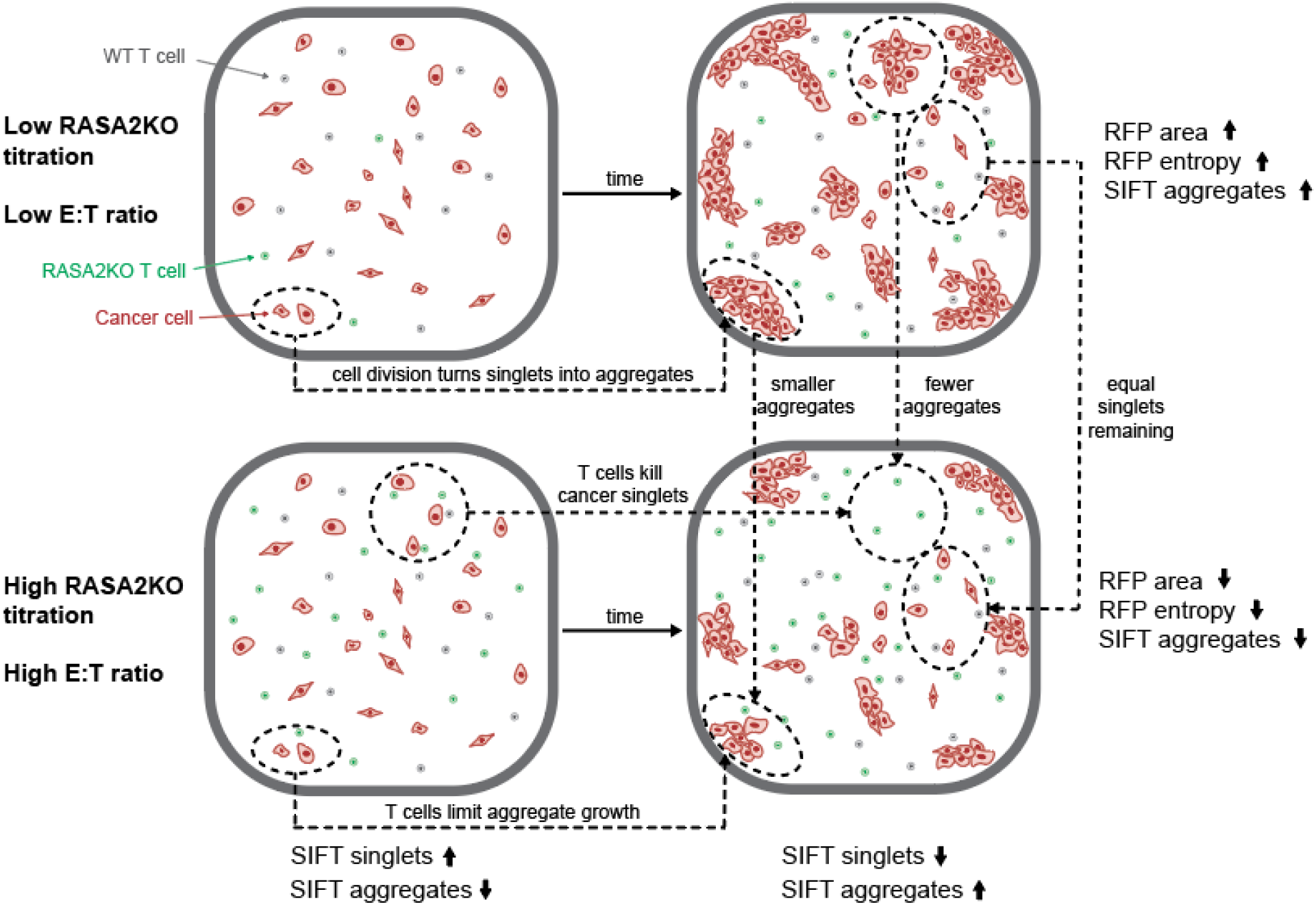
Graphical abstract of results. Schematic diagram of cancer cell aggregation dynamics from early (left column) to late time frames (right column) at low RASA2KO T cell concentrations (top row) and high RASA2KO T cell concentrations (top row). Arrow annotations along the x-axis represent SIFT singlets decreasing over time and SIFT aggregates increasing over time at both RASA2KO T cell concentrations. Arrow annotations along the y-axis represent features comparisons between the two experimental conditions at late time frames.

## Discussion

Our segmentation-free live-cell behavioral analysis (SF-LCBA) framework enabled the characterization of cancer cell aggregation dynamics at varying RASA2KO TCR T cell concentrations in low-resolution LCI co-culture experiments. By applying spatial entropy to the RFP fluorescence channel, we captured spatial patterns of cancer cell proliferation and death that extend beyond total cancer cell burden. Additionally, by clustering scale-invariant feature transform (SIFT) keypoints derived from phase images, we identified recurring local cellular topologies such as well edges, cell singlets, and multi-cellular aggregates, and we associated their abundances with distinct experimental conditions. Together, these analyses revealed that cancer cell aggregation was most effectively disrupted in co-cultures with high RASA2KO TCR T cell titrations and high effector-to-target (E:T) ratios, corroborating the findings of earlier work^13^ and validating aggregation inhibition as a potentially valuable phenotypic marker of T cell therapeutic efficacy.

While the high E:T ratio and RASA2KO titration experimental conditions were faithfully stratified by RFP area alone in our dataset, the potential of SF-LCBA lies in the orthogonal information it offers. Whereas RFP area measures total cancer cell burden, spatial entropy and unsupervised enrichment testing of SIFT keypoints reflect the spatial organization of cancer cells and their morphological state–particularly the degree of aggregation versus dispersion. In solid tumor contexts, where immune evasion is often mediated by the formation of dense tumor cell aggregates^48^, assessing the ability of T cell therapies to infiltrate and dismantle these evasive cancer cell aggregate structures is highly relevant. Our results suggest that SF-LCBA metrics may become a critical readout in future studies designed to differentiate T cell products that enhance elimination of cancer cell aggregates from those that merely clear cancer cell singlets more effectively.

This study has several important limitations. Chief among them is the relatively low spatial resolution of our LCI data (4.975 *μ*m/pixel), which is near the lower limit for detecting individual T cells (typically 5–10 *μ*m in diameter^49^). As a result, most SIFT keypoints detected in singlet-rich images were centered on cancer cells (typically 10–20 *μ*m in diameter^50^), with very few SIFT cluster singlet keypoints being centered on T cells. Additionally, low image resolution likely hindered our ability to identify mitotic cells as distinct keypoint clusters. We believe that applying SF-LCBA to images of ≥10x magnification would allow for more detailed description cancer and T cell morphologies, their patterns of co-localization, and, consequently, their phenotypic cell states. Other limitations stem from the SIFT algorithm itself. SIFT was originally designed for robust image registration by placing keypoints at high-contrast regions of the image. Consequently, many SIFT keypoints are located at the highest contrast regions of cells, most notably their edges, and therefore fails to capture multi-cellular interactions involving more than two cells. Future work where descriptor vectors, or analogous image embeddings, represent image patches spanning tens of microns could enable the detection of more complex spatial structures, such as immune infiltration gradients within aggregates. SIFT is also limited by only allowing for single-channel image inputs. While we passed phase images into SIFT in this study, future models could ingest multi-channel image data directly, enabling embeddings that better reflect local cell type composition and activity.

Despite these limitations, SF-LCBA opens up new opportunities for analyzing complex co-culture datasets in which traditional cell segmentation and tracking are infeasible. Our approach could be especially powerful in future experiments involving more complex microenvironments, including three or more interacting cell types. For example, adding dendritic cells or macrophages to T cell–cancer cell co-cultures would better recapitulate the tumor microenvironment^48^ and allow for perturbation studies across multiple immune cell types, but segmentation in these complex cell cultures is infeasible due to their high cell density and prevalence of multi-cellular interactions. Similarly, the segmentation-free nature of SF-LCBA means it could also be extended from adherent cell lines shown in this manuscript to cancer cell suspensions and patient-derived organoids, further broadening the translational potential^20,21^. Furthermore, incorporating additional fluorescent markers, such as cell viability stains, cell cycle reporters, or CD4/CD8 labels, would greatly enhance interpretability of keypoint clusters and support more nuanced phenotypic classifications^44,51,52^. Notably, even sparse use of additional fluorescent channels would suffice, as SIFT embeddings are computed from the phase channel and require fluorescence data only for cluster annotation. Collectively, the methodological improvements presented in this paper move the field closer to an ideal unsupervised framework for behavioral phenotyping in complex, multicellular systems.

## Data availability

All processed data is contained in one AnnData object that has the SIFT descriptor matrix and all the associated keypoint-, image-, and well-level metadata (including spatial entropy statistics). This data object can be freely downloaded from Zenodo at the following DOI: https://doi.org/10.5281/zenodo.15277488. Raw images will be uploaded to Bioimage Archive upon peer-reviewed publication.

## Code availability

The SF-LCBA software package, along with accompanying code to reproduce the figures shown in this manuscript, is publicly available at https://github.com/bee-hive/sflcba. Instructions for installation and usage can be found in the repository’s README.md file.

## Acknowledgements

The Marson laboratory has received research support from the Parker Institute for Cancer Immunotherapy, the Emerson Collective, Arc Institute, Sanofi, GlaxoSmithKline, and Gilead and reagents from Genscript and Illumina. L.E., A.W., and B.E.E. were funded in part by grants from the Parker Institute for Cancer Immunology (PICI), the Chan-Zuckerberg Institute (CZI), NIH NHGRI R01 HG012967, and NIH NHGRI R01 HG013736. B.E.E. is a CIFAR Fellow in the Multiscale Human Program.

## Author contributions statement

L.E., B.E.E., A.M., J.C., and A.V. conceived the study. J.C. and A.M. generated, preserved, and shared the imaging data. L.E., A.C.W., and M.S. conducted the computational experiments. L.E., A.C.W., B.E.E., J.C., and A.M. analyzed the results. L.E., A.C.W., and B.E.E. wrote the manuscript. All authors reviewed the manuscript.

## Competing interests

A.M. is a cofounder of Site Tx, Arsenal Biosciences, and Survey Genomics, serves on the boards of directors at Site Tx, and Survey Genomics, is a member of the scientific advisory boards of Site Tx, Arsenal Biosciences, Cellanome, Spotlight Therapeutics, Survey Genomics, NewLimit, Amgen, and Tenaya, owns stock in Arsenal Biosciences, Site Tx, Cellanome, Spotlight Therapeutics, NewLimit, Survey Genomics, Tenaya and Lightcast and has received fees from Site Tx, Arsenal Biosciences, Cellanome, Spotlight Therapeutics, NewLimit, Abbvie, Gilead, 23andMe, PACT Pharma, Tenaya, Lightcast, Vertex, Merck, Amgen, GLG, ClearView Healthcare, and AlphaSights. A.M. is an investor in and informal advisor to Offline Ventures and a client of EPIQ. J.C. was supported by NIH/NCI K08, 11271K08CA252605-01, a Burroughs Wellcome Fund Career Award for Medical Scientists, the Lydia Preisler Shorenstein Donor Advised Fund, and the Parker Institute for Cancer Immunotherapy, and the Pascarella Scholars Fund. B.E.E. is on the Scientific Advisory Board for Arrepath and Freenome.

## Supplementary Figures

**Figure S1.**
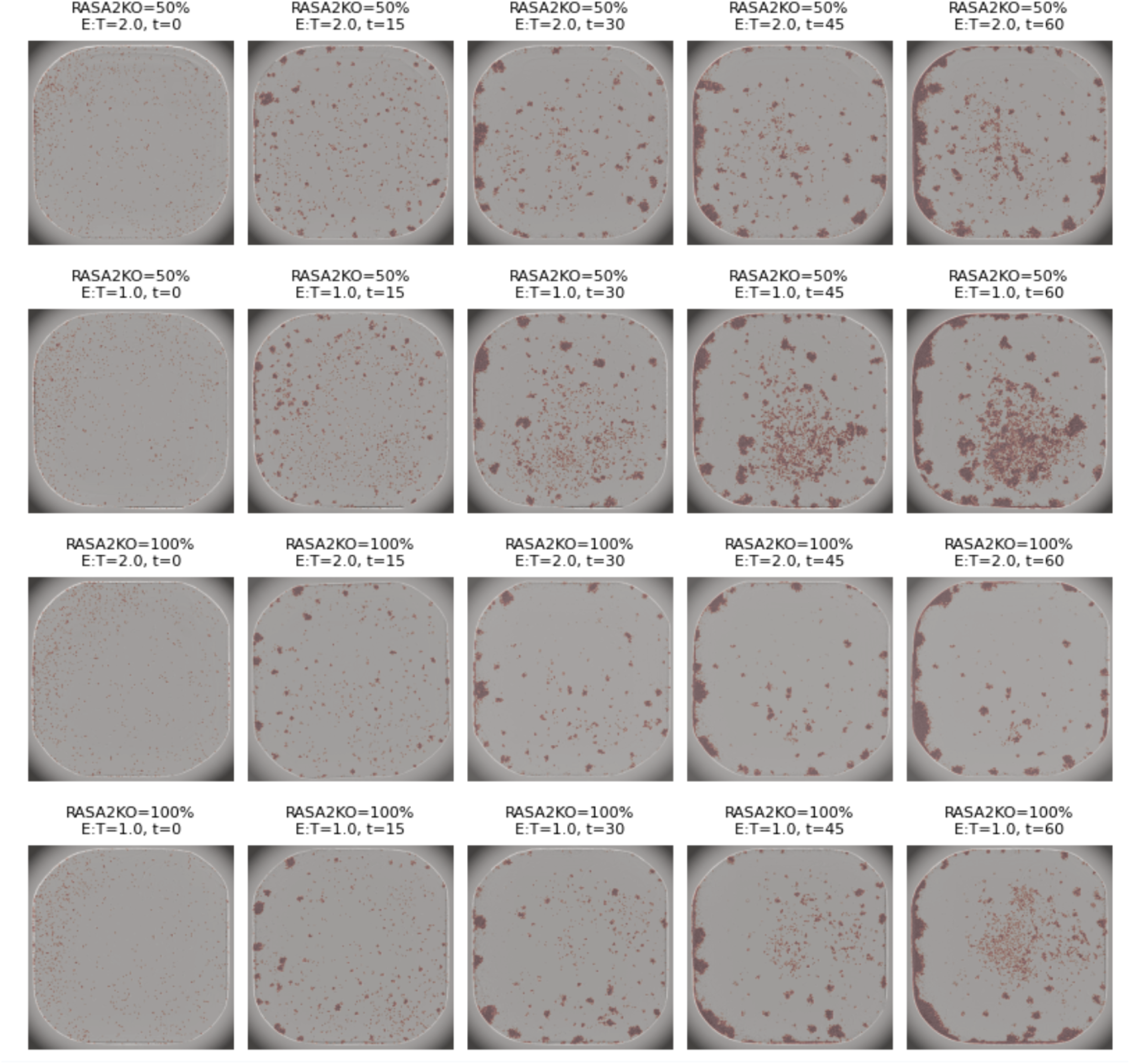
Red + phase channel images. RFP+ masks overlaid onto phase images over time at varying RASA2KO titrations and E:T ratios. Rows and columns correspond to the same images shown in Fig 1b. Time increases from left to right columns. RASA2KO titrations and E:T ratios vary between rows.

**Figure S2.**
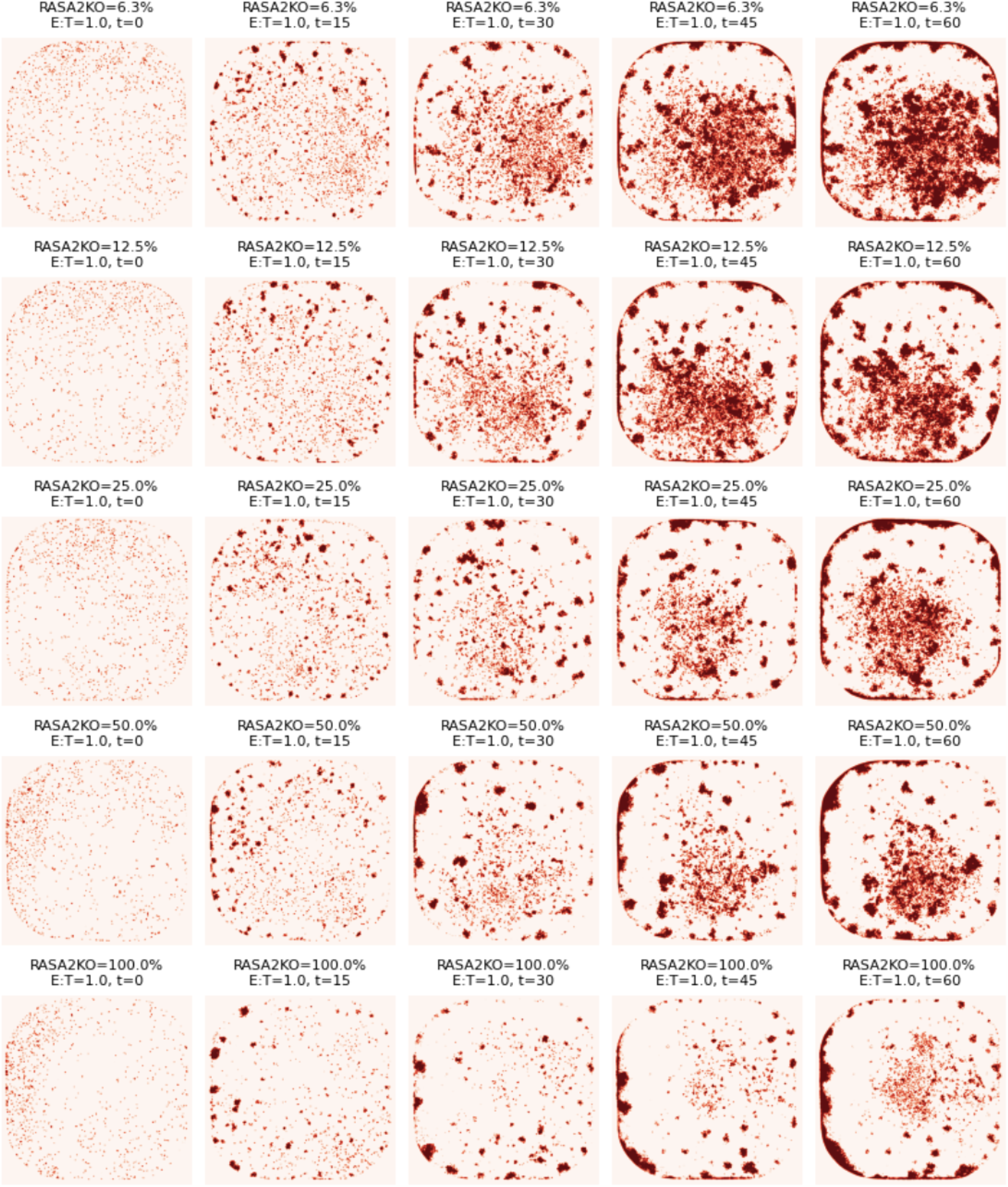
RFP+ masks vs RASA2KO titration. RFP+ masks over time at varying RASA2KO titrations and constant E:T ratio of 1.0. Time increases from left to right columns. RASA2KO titration increases from top to bottom rows.

**Figure S3.**
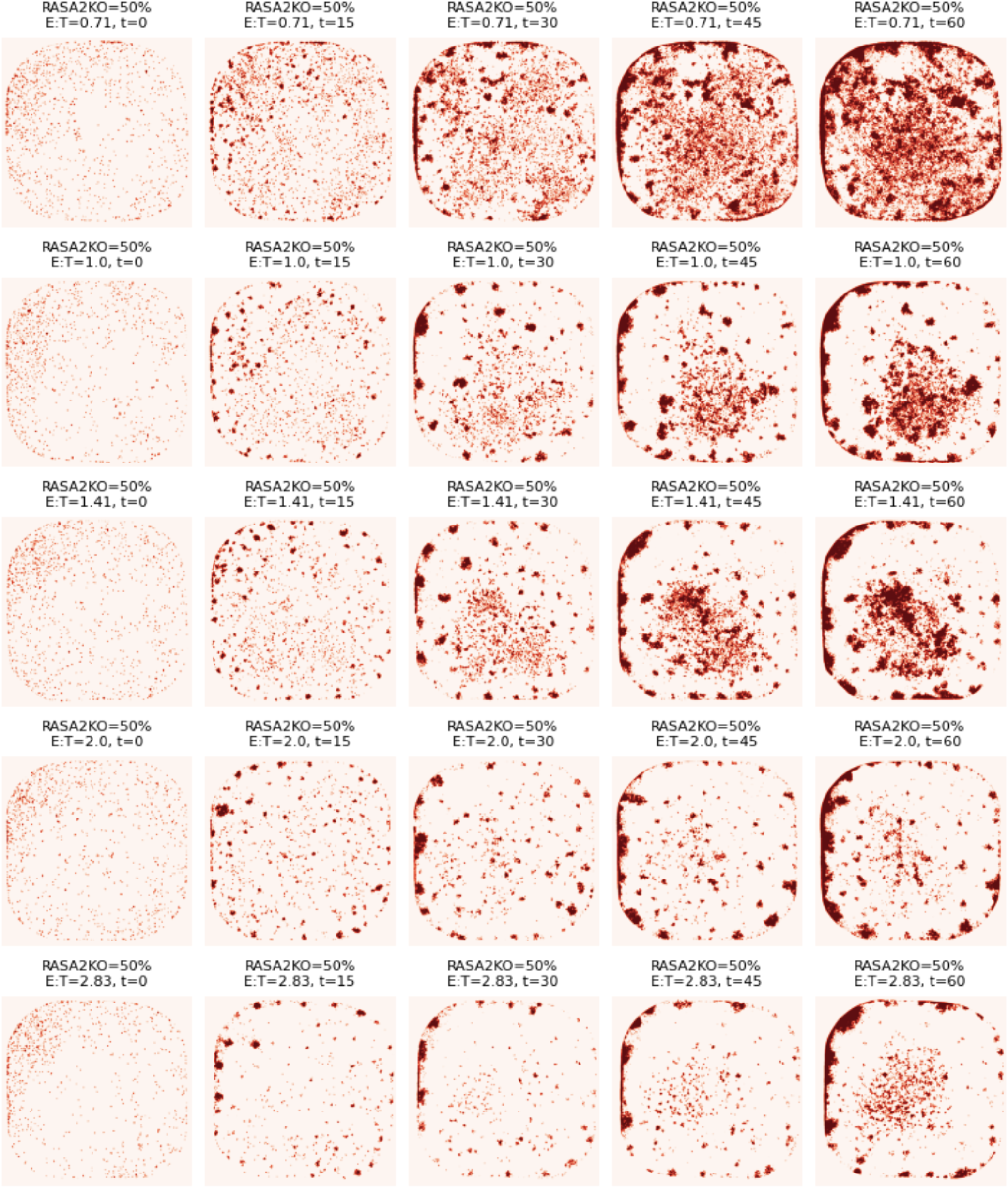
RFP+ masks vs E:T ratio. RFP+ masks over time at varying E:T ratios and constant RASA2KO titration of 50%. Time increases from left to right columns. E:T ratio increases from top to bottom rows.

**Figure S4.**
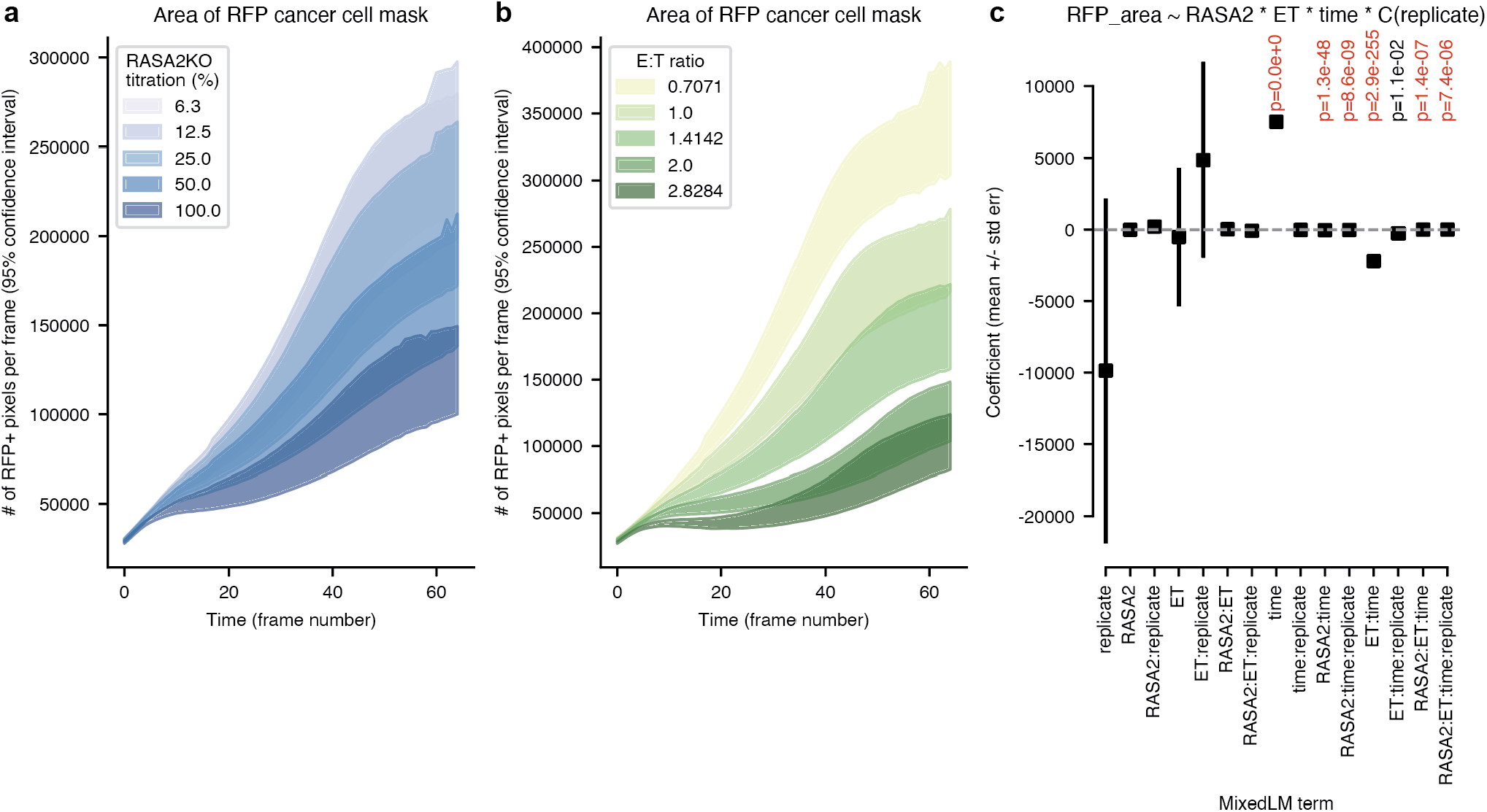
Cancer cell area dynamics. **a-b)** Cancer cell area, by measure of the number of RFP+ pixels in an image, over time, split by **a)** RASA2KO titration and **b)** E:T ratio. Shaded regions represent 95% confidence intervals. **c)** Mixed linear model results for predicting RFP+ area using the covariate values for RASA2KO titration, E:T ratio, time, and technical replicate ID, where the model is conditioned on donor ID. Donor and technical replicate are modeled as a categorical variables while the other covariates are modeled as continuous variables. The intercept term is excluded here but shown in Table 1. Boxes represent mean values of each coefficient and whiskers represent +/-standard error. Interaction terms between multiple covariates are noted using colons in the x-tick labels. Adjusted p-values (Benjamini-Hochberg correction) are annotated above each term if *p*_*adj*_ < 0.05 where the null hypothesis is that the term has no effect on RFP+ area (coefficient=0).

**Figure S5.**
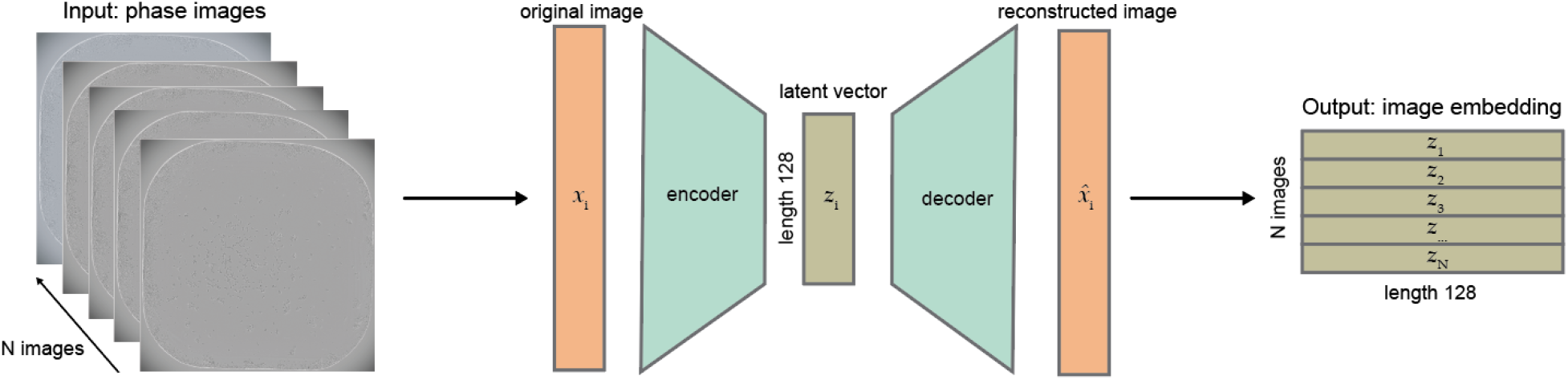
Schematic for alternative autoencoder approach. Instead of using SIFT, one could run a convolutional autoencoder on all *N* phase images in the dataset. Each image *x*_*i*_ would be passed through an encoder to produce a length-128 latent vector *z*_*i*_ which would then be passed through a decoder to reconstruct the original image 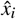. Latent vectors from all images would get concatenated to produce a final image embedding of shape *N* × 128. While the number of columns in this embedding could be set to some value other than 128 by adjusting the bottleneck size, this image embedding will always have *N* rows.

**Figure S6.**
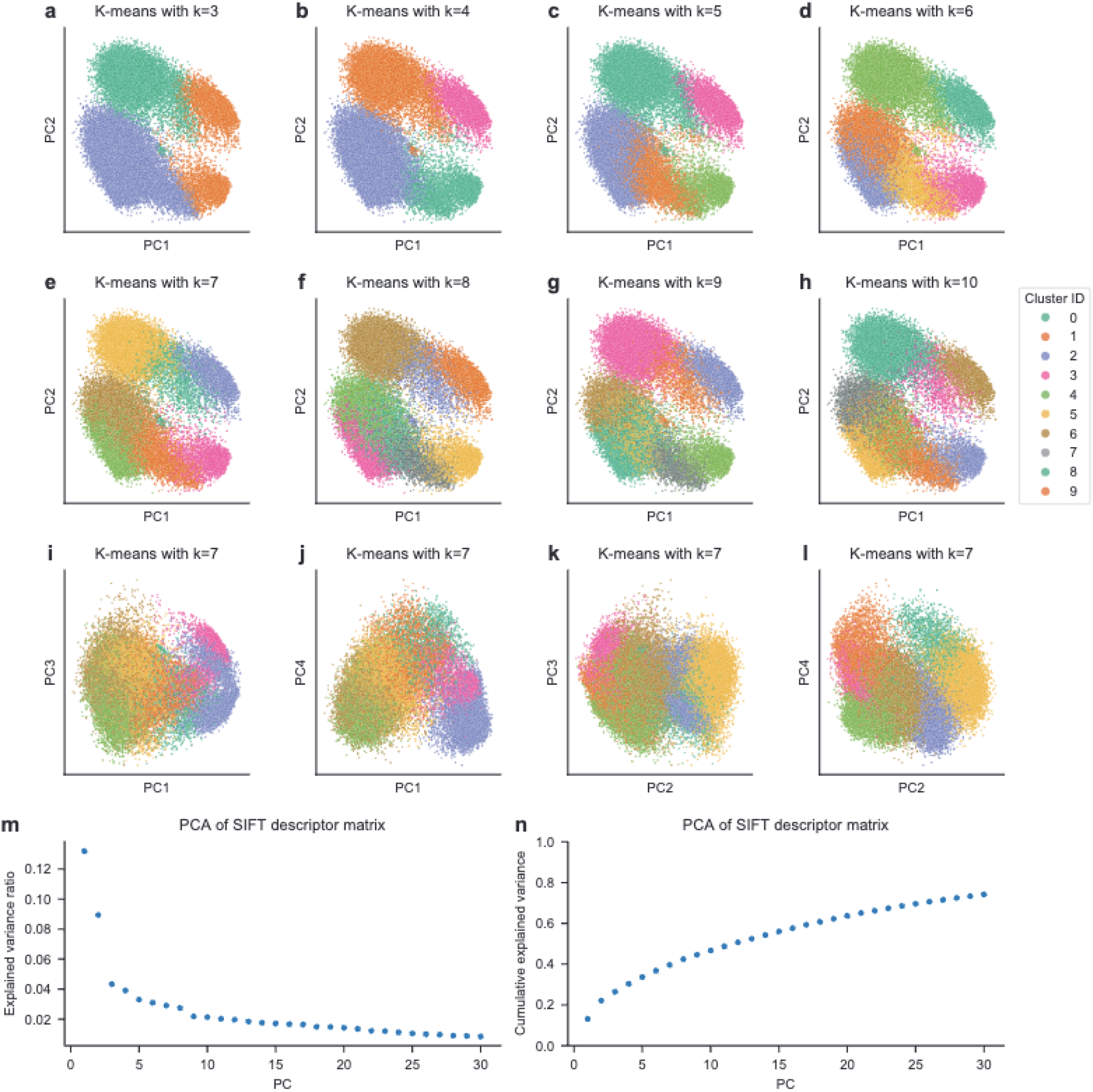
Evaluation of K-means clustering and PCA embeddings. **a-h)** First two PCs of *D* = 50, 000 SIFT descriptors colored by their K-means cluster ID. K-means clustering was performed on all *D* = 3, 389, 740 SIFT descriptors, with each subplot showing the K-means results with a different number of clusters (K) selected. **i-l)** PCA embeddings where PC3 and PC4 are shown instead of just PC1 vs PC2. Points and colors correspond to those shown in panel **e. m)** Explained variance ratio and **n)** cumulative explained variance for the top 30 PCs when running PCA on the SIFT descriptor matrix.

**Figure S7.**
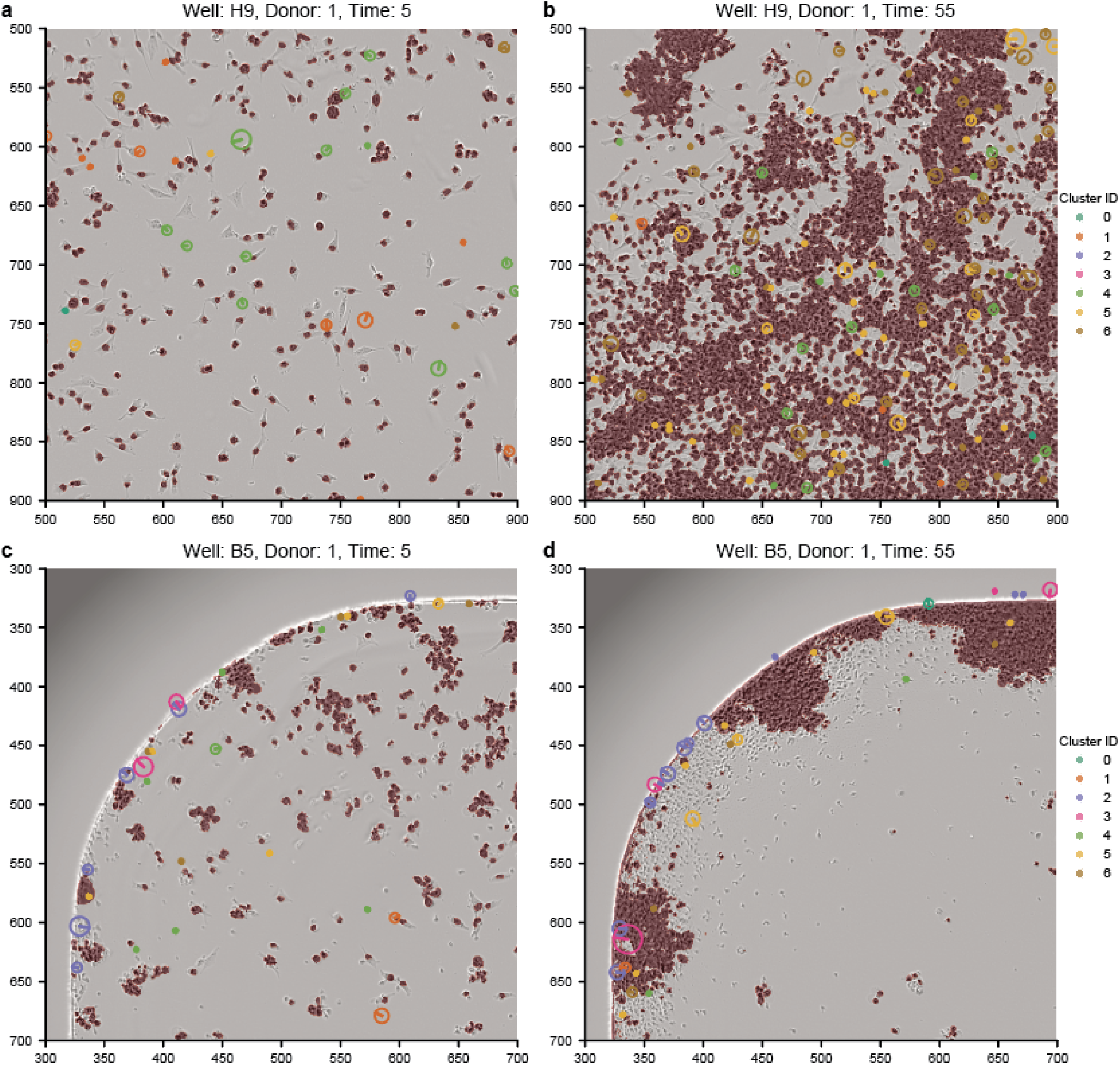
SIFT keypoints within large ROIs. Each image represents a 400 × 400 pixel ROI where the RFP mask is superimposed on top of the phase image. Colored circles are drawn around SIFT keypoints where the size corresponds to the keypoint’s radius and the color corresponds to the K-means cluster ID. Only 10% of all SIFT keypoints are shown in this image, consistent with the *D* = 3, 389, 740 subsampled dataset. **a-b)** Well H9 corresponds to RASA2KO titration of 12.5%, E:T ratio of 1.0, and ROIs located at the center of the well for time frames **a)** 5 (10 hrs) and **b)** 55 (110 hrs). **c-d)** Well B5 corresponds to RASA2KO titration of 50%, E:T ratio of 2.83, and ROIs located at the top-left corner of the well for time frames **c)** 5 (10 hrs) and **d)** 55 (110 hrs).

**Figure S8.**
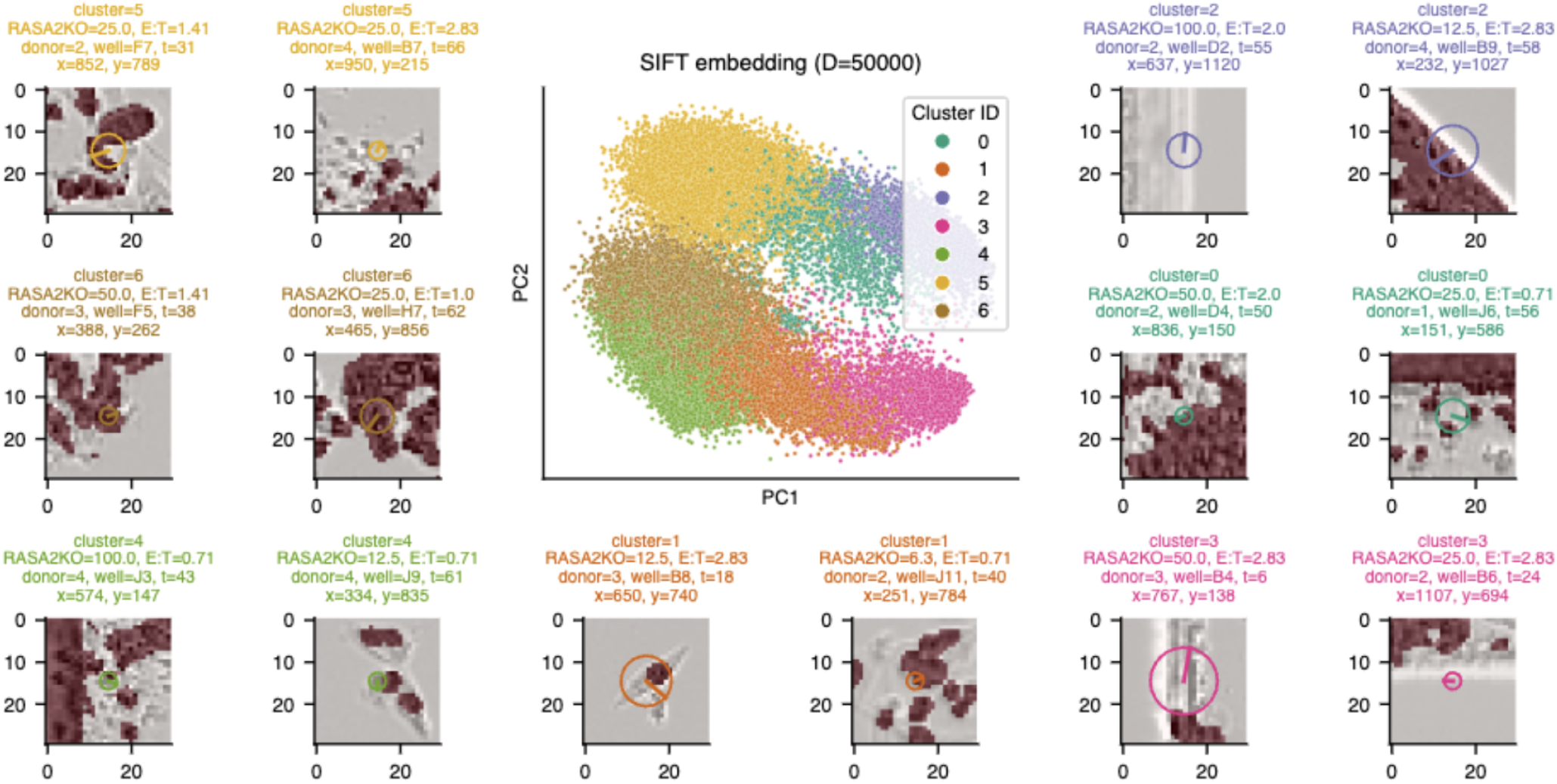
Representative ROIs for each cluster with RFP+ mask superimposed. PCA embedding of *D* = 50, 000 SIFT descriptors colored by their K-means cluster ID. PCA was run on all *D* = 3, 389, 740 SIFT descriptors prior to subsetting to *D* = 50, 000. Two representative ROI phase images are shown for each cluster with their RFP+ cancer cell masks superimposed. The circle in each ROI represents the radius of each SIFT keypoint and the line represents its relative orientation. ROIs are annotated with the SIFT keypoint’s xy-coordinate along with the associated image metadata.

**Figure S9.**
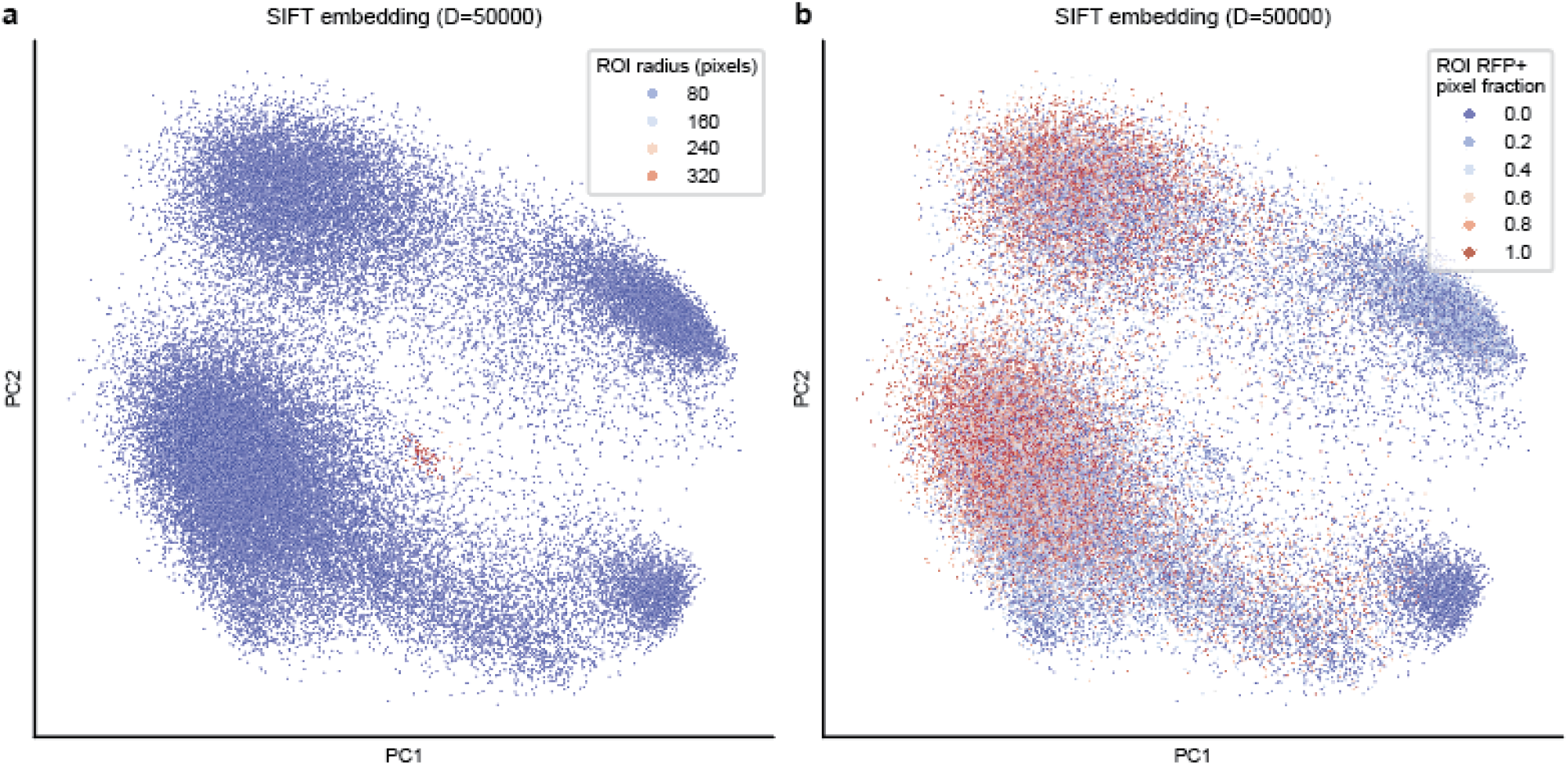
ROI properties overlayed onto SIFT embeddings. PCA embedding of *D* = 50, 000 SIFT descriptors colored by their ROI a) radius and b) fraction of RFP+ pixels. Radius *r* for each SIFT descriptor is defined by its scale *σ* and octave *o* by the equation *r* = *σ* * 2^1+*o*^.

## Notes

https://github.com/bee-hive/sflcba

